# Brain serotonin circuit reverses decline in physical activity with age

**DOI:** 10.64898/2025.12.27.695444

**Authors:** Pablo B. Martinez de Morentin, Alasdair Leeson-Payne, Shuwen Mu, Yuliia Martynova, Aida Mohammadkhani, Drew M. Neyens, J. Antonio Gonzalez, Janis Lazovskis, Claudia Cristiano, Daniel R. Crabtree, Claire L. Fyfe, William Buosi, Daniel Powell, Raffaella Chianese, Matevz Arcon, Guy S. Bewick, Arimantas Lionikas, Graham Horgan, Ran Levi, Alexandra M. Johnstone, Stephanie L. Borgland, Lora K. Heisler

## Abstract

Physical inactivity increases with age and is the fourth leading risk factor for mortality. Establishing the mechanisms underpinning declining physical activity (PA) with age has remained both elusive and difficult to overcome. Older (50-70 years old) compared with younger (17-30 years old) UK participants showed reduced PA, and this profile was effectively modelled in mice. Analysis of brain serotonin (5-HT) neurons in the dorsal raphe (DR) revealed altered firing activity in older mice. Chemogenetically mimicking this 5-HT^DR^ tone in young adult mice produced an older adult PA profile. Genetically blocking 5-HT activity at 5-HT_2C_ receptors (5-HT_2C_Rs) prevented the decline in PA with age. Importantly, barring 5-HT action at 5-HT_2C_Rs specifically within the ventral tegmental area restored youthful PA levels and strength in older mice. These data fill a longstanding knowledge gap by defining brain circuitry programming the decline in PA with age, and importantly, a means to reverse it.

## MAIN

Regular physical activity (PA) is inversely associated with all-cause mortality and the risk of numerous morbidities, including cardiovascular disease, type 2 diabetes, cancer, depression, and dementia.^1-6^ This strong relationship between PA and health has been demonstrated across a broad spectrum of studies, including randomized controlled trials,^7-10^ mechanistic investigations,^11-14^ and mammalian models.^15-18^ An early and influential study by Chapman and colleagues (1968) demonstrated the rapid and reversible impact of inactivity on health. Specifically, a three-week period of inactivity via enforced bed rest in healthy 20-year-old men produced cardiovascular and metabolic profiles consistent with men twice their age, whereas resuming PA restored their original health status.^19^ This and subsequent studies over the last 50 years clearly demonstrate the causal and dynamic relationship between PA/sedentary behavior and physiological aging.

Although regular activity is essential for maintaining both physical and mental health, habitual PA levels progressively decline with age.^20, 21^ From around the age of 30, adults lose approximately 3–8% of skeletal muscle mass per decade, alongside reductions in energy expenditure, which adversely affects metabolic health, functional independence, and quality of life.^22-24^ Midlife therefore represents a critical period to preserve youthful activity levels to prevent disease and extend healthspan.^25-27^ The 50–70-year-old age bracket is particularly important yet relatively understudied in this context.^28^ Notably, individuals within this demographic are projected to constitute one in four adults by 2038 in some countries,^29^ emphasizing the need for targeted research examining PA behavior and mechanisms in this age group.

Historically, research has primarily focused on the impact of moderate-to-vigorous physical activity (MVPA) on disease prevention and longevity. However, accumulating evidence demonstrates that even lower-intensity or incidental PA provides significant health benefits, whereas prolonged inactivity has detrimental effects.^10, 30^ Walking, the most common and accessible form of PA, is now measurable in free-living conditions through wearable technologies. Walking pace shows an inverse association with all-cause mortality, cardiovascular disease, type 2 diabetes, and cognitive decline in middle-aged and older adults.^6, 31-33^ These findings indicate that health protection is determined not only by the volume of activity but also by walking intensity and pace.^34^

While the major brain regions and neurotransmitters governing locomotor activity have been progressively elucidated over recent decades, the neuronal mechanisms underlying the age-associated decline in voluntary PA remain incompletely understood. The evolutionarily conserved neurotransmitter serotonin (5-hydroxytryptamine; 5-HT) exerts wide-ranging effects on behavior and physiology across species,^35, 36^ including an inverse relationship with locomotor activity.^37^ Increasing 5-HT bioavailability through selective serotonin reuptake inhibitors (SSRIs) reduces locomotor activity across diverse taxa, including invertebrates (e.g., *Drosophila*), aquatic vertebrates (e.g., zebrafish), and terrestrial mammals (e.g., rodents and humans).^37-39^ In older human adults, the use of SSRIs is associated with reduced exercise capacity, grip strength, and walking speed.^40^

Mechanistic studies implicate 5-HT neurons within the dorsal raphe nucleus (5-HT^DR^) in modulating locomotion. Acute chemogenetic and optogenetic activation of 5-HT^DR^ neurons suppresses exploratory behavior in the open field and elevated plus maze tests.^41-43^ However, given the potential confounding influence of anxiety in these assays, it remains unclear whether these effects reflect a direct suppression of locomotion *per se* or indicate changes in anxiety-associated behavior. To address this, we investigated how selective manipulation of 5-HT^DR^ neuronal activity alters spontaneous home cage PA parameters.

In mammals, 5-HT mediates its diverse effects through at least 14 receptor subtypes, each with distinct distribution and function.^44^ Converging pharmacological and genetic evidence identifies the G_q_-coupled 5-HT_2C_R receptor (encoded by the *Htr2cr* gene) as a primary mediator of 5-HT’s inhibitory effects on locomotion. Activation of 5-HT_2C_Rs reduces ambulation, while receptor antagonism or genetic inactivation enhances activity levels.^45-47^ 5-HT_2C_Rs are expressed in the brain,^48^ however, the specific neuronal populations mediating its control of PA remain unclear. 5-HT_2C_R agonists inhibit mesolimbic dopamine (DA) activity and release,^49-51^ suggesting a potential 5-HTergic modulation of DAergic circuits involved in motivation and movement. Activation of DA neurons within the ventral tegmental area (VTA) robustly enhances spontaneous PA in rats.^52^ We therefore investigated the specific role of 5-HT_2C_R activity within the VTA in regulating DA neuronal firing and age-related declines in voluntary PA.

Given that PA is a key modifiable risk factor for chronic disease and premature mortality, this study addressed several critical questions: (i) To what extent is the age-related decline in PA biologically conserved across species?; (ii) Is the modulation of 5-HT signaling sufficient to mimic or reverse age-associated PA profiles?; (iii) Does chronic blockade of 5-HT_2C_Rs prevent age-related decline in voluntary PA?; and (iv) Can selective ablation of 5-HT_2C_R^VTA^-expressing neurons restore youthful PA patterns in older mice? Collectively, these investigations address a fundamental question in the biology of aging: through which mechanisms does voluntary PA progressively decline with age and can this process be reversed?

## RESULTS

### Cross species age-related decline in PA

Here we investigated the extent to which changes in PA are biologically mediated by comparing the human age-related PA profile with that of mice housed in a constant environment and conditions throughout adulthood. To assess adult PA levels, we analyzed data collected from female (n=45) and male (n=29) Scottish participants from the Full4Heath study.^53^ Younger adults (17-30 years old) and older adults (50-70 years old) were provided with accelerometers for one week, and average daily caloric intake was assessed with the 24 hr multiple pass dietary method on four occasions. Body weight and composition were assessed under controlled conditions at school or the Human Intervention Studies Unit at the Rowett Institute, Scotland (**Table 1**). The analysis revealed that older adults (aged 50-70) showed less PA, measured both as daily distance (**Figure 1A**) and a slower speed compared to younger adults (aged 17-30; **Figure 1B**). Percentage of time within a gait speed category was also examined. This revealed that the older adult group spent more time within a sedentary/slow cadence (0-30 steps per minute) and less time engaged in a moderate walking speed (60-90 steps per minute), brisk walking (90-120 steps per minute) or fast ambulatory activity (120-240 steps per minute) classification compared with younger adults (**Table 1**). Older participants also had a higher body mass index (BMI; **Figure 1C**), higher adiposity as percent body fat (**Figure 1D**) and lower lean body mass (**Figure 1E**). We observed no significant differences in self-reported daily caloric intake between the two groups of adults (**Figure 1F**).

**Figure 1.**
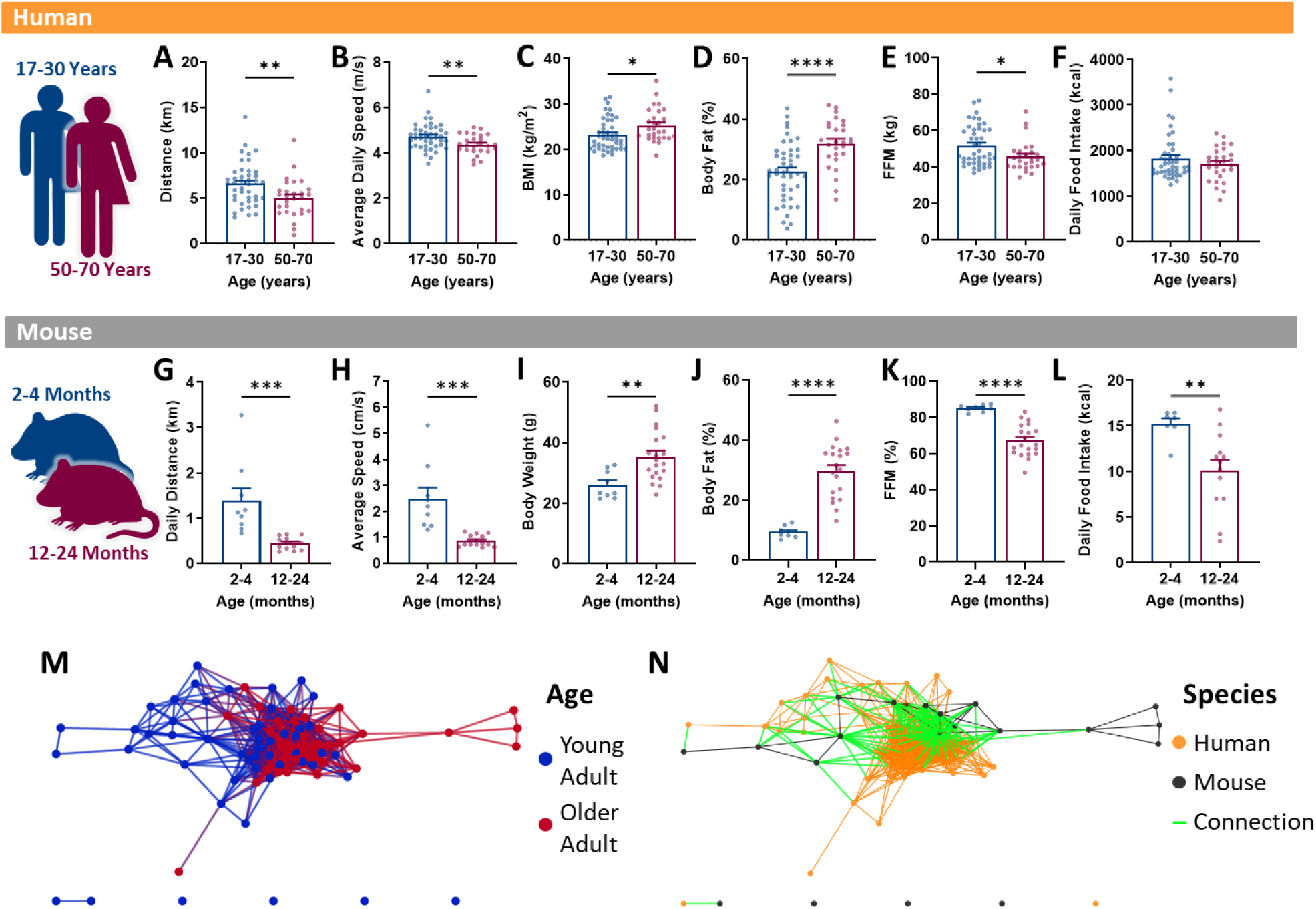
Age-related decline in human physical activity parameters is modelled in mice. **A-F.** Compared to younger adults aged 17-30 (n=45), older adult male and female Scottish volunteers aged 50-70 (n=29) show: (A) reduced average daily distance (t(68) = 2.817, p = 0.0063); (B) lower average daily speed (t(68) = 2.698, p = 0.0088); (C) higher body mass index (BMI) (t(70) = 2.343, p = 0.0220); (D) higher body fat (t(69) = 4.160, p = <0.0001); (E) lower fat free mass (t(69) = 2.422, p = 0.0181); (F) no change in daily food intake (t(70) = 1.089, p = 0.2797). **G-L.** Compared to younger adult (n=9) male and female mice, older adult (n=20) mice exhibit: (G) reduced average daily distance (t(20) = 4.092, p = 0.0006); (H) lower average daily speed (t(20) = 4.474, p = 0.0002); (I) higher body weight (t(27) = 3.149, p = 0.0040); (J) higher body fat (t(27) = 6.711, p = <0.0001); (K) lower fat free mass (t(27) = 5.994, p = <0.0001); (L) lower daily caloric intake (t(18) = 2.976, p = 0.0081). **M-N.** Geographic graphing of 6-variables (A-F or G-L) in 6-dimensional space creating a sphere for each subject, clustered and linked with connecting lines based on similarity shows: (M) younger adult participants and mice (blue spheres and lines) have a high degree of similarity illustrated by clustering left compared to older adults (red spheres and lines) shown clustering right. (N) intermixed species, with most mice (black) showing a high degree of similarity (green connecting lines) in their metabolic signature to humans (orange).

Human life phase equivalent female and male mice housed in consistent conditions during adulthood were used to measure natural, home cage PA patterns (younger adult: 2-4 months old and older adult: 12-24 months old). PA parameters and food intake levels were noninvasively and automatically measured over one week. Consistent with the older adult human participants, older adult mice were significantly less active (**Figure 1G**), had a lower average ambulation speed (**Figure 1H**), higher body weight (**Figure 1I**), higher percent body fat (**Figure 1J**) and lower lean mass (**Figure 1K**). In contrast to the human study, we found that younger adult mice consumed more daily calories compared with older adult mice (**Figure 1L**).

A limitation of conventional data graphing is that individuals are represented as unlinked data points between the bar graphs (e.g. **Figure 1A-F and 1G-L**), obscuring the composite profile of each individual. Geographic graphing methods combine normalized data for each human participant or mouse by creating an amalgamation of linked data points from each of these 6 variables in 6-dimensional space, represented as a sphere. This composite provides an individualized ‘metabolic signature’ for each human participant and mouse. To visualize similarities and differences between the subject’s data spheres, a radius threshold of 24 units around each subject was selected, with a distance measured as Euclidean distance to show how each subject is related to all other subjects. Each dot in **Figure 1M** represents a composite sphere (metabolic signature) for each human participant or mouse. If the spheres between subjects intersect (denoting similarity), then an edge connecting the two subjects is displayed. Interpreting the metabolic data in 6-dimensional space reveals that the older subjects show a similar metabolic signature that cluster them together (e.g. red branches clustering to the right), and the same for younger participants and mice (e.g. blue branches clustering to the left; **Figure 1M**). Changing the colors to visualize species of these subject spheres reveals direct similarity between humans and mice (**Figure 1N**). Specifically, most mice (black spheres) showed a direct connection (green line) with human participants (orange spheres). Taken together, our data illustrate that humans and mice share a strikingly similar aging-related multi-characteristic profile. We therefore concluded that mice model the biological decline in PA with aging observed in people, and mice may be used to interrogate the neural underpinnings driving waning PA with age.

### 5-HT tone is inversely related to PA and is altered in older adult mice

To elucidate the mechanisms through which voluntary PA is controlled by the brain, we turned our attention to the neurotransmitter 5-HT, levels of which have an inverse relationship with locomotor output. We first examined whether an older PA profile could be generated in young adult mice by increasing 5-HT bioavailability. To achieve this, young adult wild type mice were housed in TSE PhenoMaster chambers that non-invasively collect PA patterns. Prior to the onset of the dark cycle, mice were treated with d-fenfluramine, a drug which promotes the release of endogenous 5-HT and blocks its reuptake (**Figure 2A**). Consistent with data obtained in rats,^54^ we observed that d-fenfluramine (3 mg/kg, i.p.) significantly reduced PA levels. Specifically, d-fenfluramine decreased dark cycle home cage activity illustrated by a heat map visualizing total ambulation within the cage zones (**Figure 2B**) and as quantified by hourly horizontal beam breaks (**Figure 2C**). In addition to reducing activity, d-fenfluramine also increased the percentage of time mice spent inactive compared to vehicle treatment (**Figure 2D**). A more detailed analysis of activity patterns revealed that d-fenfluramine significantly reduced the amount of daily ambulation measured as the distance mice travelled within the home cage (**Figure 2E**) and their average ambulation speed (**Figure 2F**) compared with vehicle treated mice. Mice administered with d-fenfluramine at this dose showed no changes in food intake (**Figure 2G**). These data reveal that broadly increasing endogenous 5-HT availability in young adult mice rapidly produces a PA profile consistent with older adult mice (**Figure S1**).

**Figure 2:**
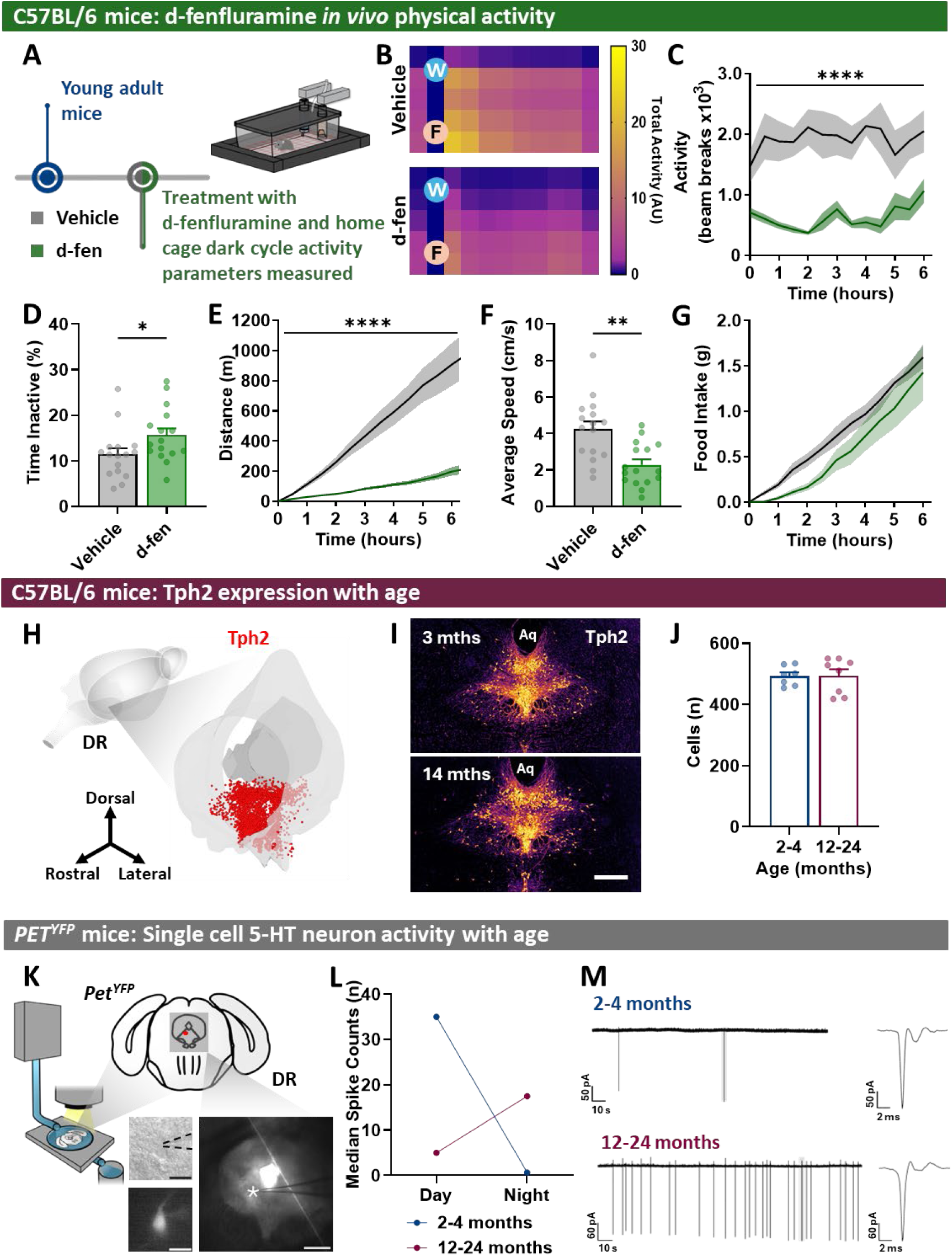
Older adult physical activity phenotype is associated with increased 5-HT^DR^ neuron activity. **A-G.** (A) Effect of d-fenfluramine (d-fen, 3 mg/kg, i.p.) versus vehicle (saline) on TSE PhenoMaster home cage physical activity in young adult (2-3 months old) male and female mice on a C57BL/6 background (n = 16) showed: (B) reduced beam breaks as illustrated in representative ambulation home cage heatmaps (W, water; F, food); (C) decreased total beam breaks over time (RM ANOVA Treatment: F_(1,14)_ = 37.13, p = <0.0001); (D) increased percent time spent inactive (t(30)= 2.101, p = 0.0442); (E) reduced cumulative daily distance (RM ANOVA Treatment: F_(1,14)_ = 30.72, p = <0.0001); (F) decreased average speed (t(15) = 3.467, p = 0.0034); (G) no change in cumulative food intake (RM ANOVA Treatment: F_(1,14)_ = 2.542, p = 0.1332). **H-J.** Dorsal raphe nucleus 5-HT expression. (H) Spatial representation of 5-HT^DR^ neurons using Tph2 immunoreactivity in a young adult mouse. (I) Representative heatmap of Tph2 positive cells expression in DR at bregma -4.60 (scale bar = 400 µm). (J) Tph2-immunofluroescent cell number at bregma level -4.60 (2-4 months old, n = 7; 12-24 months old, n=8; t(13) = 0.04933, p = 0.9614). **K-M**. (K) 5-HT^DR^ cell activity in younger (2-4 months old, n=4 mice/36 cells) and older (12-24 months old, n=4/20 cells) adult male and female *ePet^YFP^* mice in the light and dark cycle (scale bars: main image = 500 µm; magnified = 20 µm). (L) Median spike count demonstrates 5-HT^DR^ neurons from older adult mice exhibit increased firing in the dark cycle and lower firing in the light cycle compared to younger adult mice (ANOVA Interaction F_(1,52)_ = 7.433, p = 0.0087). (M) Example traces during the dark cycle.

To establish whether the endogenous activity of 5-HT neurons changes with age and whether this could explain the age-related reduction in PA, we profiled 5-HT neurons in younger and older adult mice. We focused on the neuronal population forming the principal source of 5-HT in the brain, the DR, and plotted its expression in three-dimensions (**Figure 2H**)^44^. Immunohistochemistry used to visualize 5-HT cells (tryptophan hydroxylase 2, Tph2+) revealed no differences in the number of 5-HT^DR^ neurons between younger and older adult mice (**Figure 2I-J**). Next, cell-attached patch-clamp electrophysiology was employed in *ePet-yellow fluorescent protein* (*ePet^YFP^*) mice to compare the activity profile of 5-HT^DR^ neurons during the light and dark cycle (n=56 cells; **Figure 2K**). We observed a significant age interaction, with 5-HT^DR^ neurons from older adult *ePet^YFP^* mice showing greater activity during the dark cycle and less activity during the light cycle compared to younger adult mice (**Figure 2L-M**). Given that increasing 5-HT bioavailability is associated with reduced locomotion, greater endogenous 5-HT^DR^ neuron activity in older adult mice during their primary wakeful period (dark cycle) could contribute to their lower PA profile.

### Increased 5-HT^DR^ neuron activity promotes an older-age PA signature in young mice

We next probed whether mimicking older adult increased dark cycle 5-HT^DR^ neuron activity would be sufficient to create an older adult mouse PA profile in young adult mice *in vivo*. Chemogenetic Designer Receptors Exclusively Activated by Designer Drugs (DREADD) was employed to control 5-HT^DR^ cellular activity. Adeno-associated viruses (AAVs) expressing h3MDq-mCherry (AAV8-hSyn-DIO-hM3D(Gq)-mCherry; 5-HT^DR^:hM3Dq) or control mCherry (AAV8-hSyn-mCherry; 5-HT^DR^:mCherry) were delivered into the DR of young adult *Tph2^iCre^* mice (**Figure 3A-B**). Administration of designer drug clozapine-N-oxide (CNO, 1 mg/kg, i.p.) to young adult 5-HT^DR^:hM4Dq mice to increase 5-HT^DR^ activity was sufficient to produce an older adult PA profile. Specifically, CNO significantly reduced dark cycle home cage PA as visualized with heat maps (**Figure 3C**) and measured with horizontal beam breaks over time (**Figure 3D**). CNO also increased time inactive (**Figure 3E**). Assessment of ambulation revealed CNO decreased distance travelled (**Figure 3F**) and average speed (**Figure 3G**) compared with vehicle administration. As expected, CNO treatment had no effect in control 5-HT^DR^:mCherry mice (**Figure S2**). These data illustrate that increasing 5-HT^DR^ neuron activity promotes an older age PA phenotype in young adult mice. This result was specific to PA and did not alter another behavior central to 5-HTergic signaling, namely food intake (**Figure 3H**).

**Figure 3:**
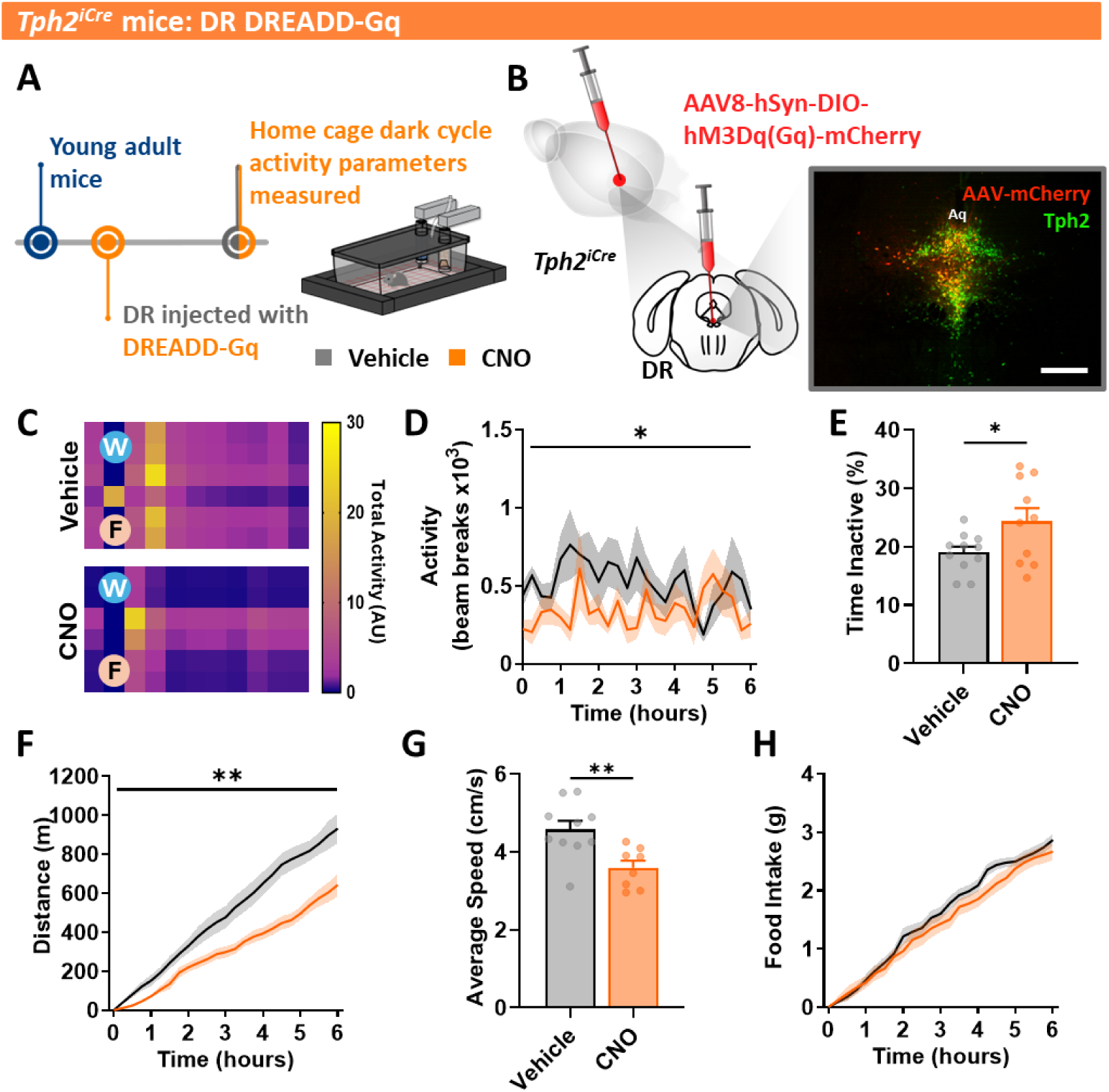
Increased 5-HT^DR^ neuron activity in young adult mice promotes an older adult physical activity signature. (A) Schematic of chemogenetic study design in young adult *Tph2^iCre^* mice (4 months old, n=11). (B) Schematic of AAV delivery into the DR and representative immunofluorescence image of hM3Dq(Gq)-mCherry (red) and TPH2 neurons (green) (Aq: cerebral aqueduct; scale bar = 400 µm). **C-H.** Compared to vehicle (saline) control, treatment with clozapine-n-oxide (CNO; 1 mg/kg, i.p) rapidly promoted a profile consistent with older mice in young adult mice as shown by: (C) reduced representative ambulation home cage heatmaps (W, water; F, food); (D) reduced total home cage activity (RM ANOVA Treatment: F_(1,19)_ = 5.815, p = 0.0262); (E) increased time inactive (t(19) = 2.214, p = 0.0392); (F) reduced distance travelled (RM ANOVA Treatment: F_(1,18)_ = 12.86, p = 0.0021); (G) decreased average ambulation speed (t(16) = 3.249, p = 0.0050); (H) no alteration in cumulative food intake (RM ANOVA Treatment: F_(1,19)_ = 0.9944, p = 0.3312).

### Prevention of age-related decline in PA in *5-HT_2C_R* knockout mice

The 5-HT receptor with the strongest link to PA as evidenced by pharmacological and murine gene knockout studies is the 5-HT_2C_R.^45-47, 49-51^ We investigated whether abolishing 5-HT action exclusively at this receptor is sufficient to prevent the reduction in PA with aging (**Figure 4A**). Consistent with this hypothesis, older adult *loxTB Htr2cr* knockout (*5-HT_2C_R*^KO^) mice were protected from the age-related decline in PA. Specifically, as visualized in heat maps of home cage ambulatory activity (**Figure 4B**) and quantified via beam breaks over time (**Figure 4C**), older adult *5-HT_2C_R*^KO^ mice exhibited a pattern of PA consistent with young adult mice (**Figure S1**). Like young adult mice, older adult *5-HT_2C_R*^KO^ mice spent less time inactive as compared with their wild type siblings (**Figure 4D**). Daily distance travelled (**Figure 4E**) and average speed (**Figure 4F**) were comparable to young adult mice PA (**Figure S1**). Food intake was increased near the onset of the dark cycle, but not later in the dark cycle (**Figure 4G**). These data signify that blocking age-related increased 5-HT^DR^ activity at the 5-HT_2C_Rs is sufficient to prevent the reduction in PA with aging.

**Figure 4:**
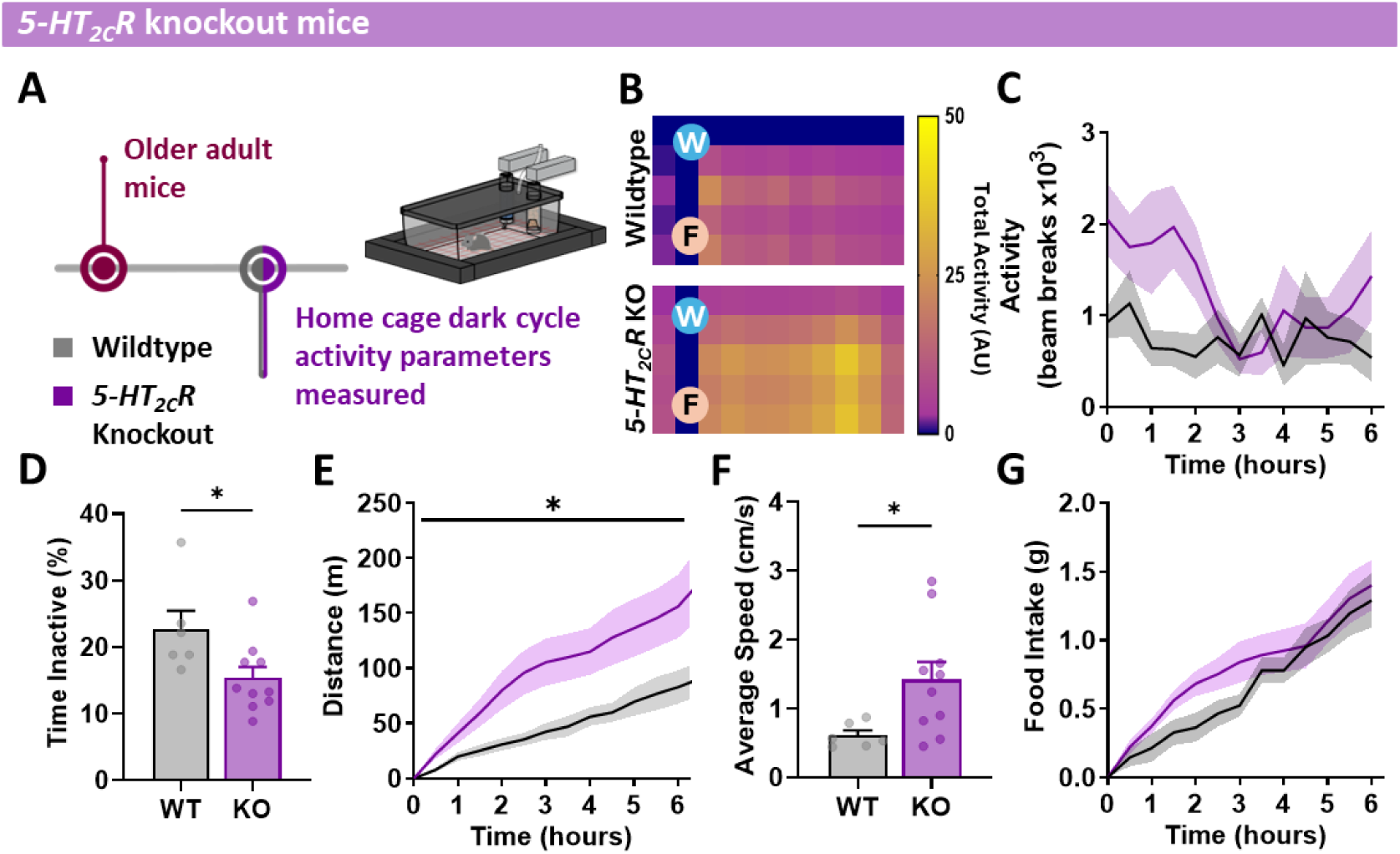
*5-HT_2C_R^KO^* prevents decline in physical activity with age. (A) Schematic detailing study design of physical activity assessment of older adult (12-16 months old) *5-HT_2C_R* knockout (KO, n=8) and wildtype (WT, n=6) mice. *5-HT2CR*^KO^ mice are protected from the decline in physical activity with age as illustrated by: (B) representative home cage beam break heatmaps (W, water; F, food); (C) increased home cage total activity during the first 6 hours of the dark cycle (RM ANOVA Genotype: F_(1,14)_ = 4.525, p = 0.0517); (D) reduced time inactive (t(14) = 2.393, p = 0.0313); (E) greater distance travelled (RM ANOVA Genotype: F_(1,12)_ = 5.314, p = 0.0398); (F) faster average speed (t(14) = 2.359, p = 0.0334); (G) no genotypic effects on cumulative food intake (RM ANOVA Genotype: F_(1,14)_ = 1.128, p = 0.3063).

### 5-HT2CRs indirectly inhibit DA^VTA^ neurons via activation of GABAergic interneurons

We examined the neural underpinnings of this 5-HT → 5-HT_2C_R PA modulation. Brain 5-HT inhibits locomotion in part via reciprocity with the DAergic system, a neurotransmitter modulating motivation, reward and physical activity. 5-HT^DR^ neurons innervate the VTA, a region expressing dopamine neurons (DA^VTA^)^55-57^ which control PA^52^ and a particular abundance of 5-HT_2C_Rs^58^. We began by visualizing the 5-HT^DR^ ◊ VTA circuit by injecting mice with the retrograde canine adenovirus expressing green fluorescent protein (CAV2-GFP) that labels axons and somas into the VTA. We observed that 5-HT neurons within the bregma level of the DR that we targeted using chemogenetics (**Figure 3B**) innervate the VTA (**Figure 5A-B**). 5-HT released into the VTA acting at 5-HT_2C_Rs could either directly, or indirectly via GABAergic interneurons (VTA^GABA^), impact the activity of DA^VTA^ neurons. Immunohistochemical staining for 5-HT_2C_Rs, GABA (using *vgat-IRES-Cre:Rosa-tdTomato*; *vGat^Cre:tdTom^* mice; a marker for GABA neurons) and DA (using tyrosine hydroxylase, TH; a marker for dopamine neurons) revealed that 5-HT_2C_Rs are primarily co-expressed with GABA^VTA^ neurons, not DA^VTA^ neurons (**Figure 5C-D**). These findings are consistent with previous reports using different methods.^58, 59^ 5-HT_2C_Rs are situated to modulate DA activity through more than one mechanism. However, based on our own and other earlier work,^51, 58^ we hypothesized that a primary effect of 5-HT is to indirectly inhibit DA^VTA^ neurons via action at 5-HT_2C_Rs in GABA^VTA^ interneurons (**Figure 5E**).

**Figure 5.**
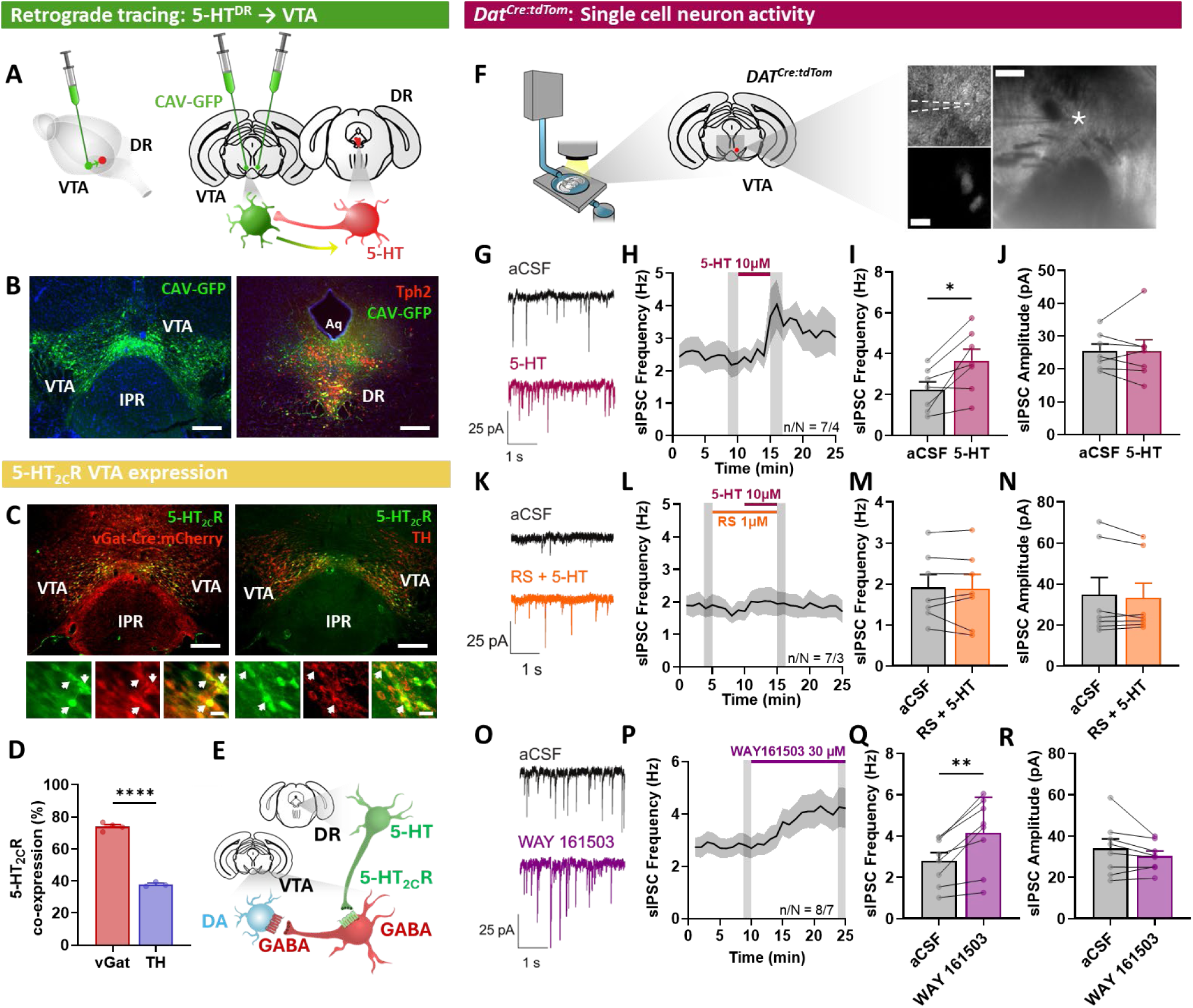
5-HT^DR^ neurons indirectly inhibit DA^VTA^ neurons via activation of 5-HT_2C_R^VTA^ GABAergic interneurons. (A) Retrogradely transported CAV-GFP injected into the VTA to establish the extent and localization of 5-HT^DR^ neurons projecting to the VTA. (B) Representative immunofluorescence images of CAV-GFP (green) injection into the VTA (left panel) and 5-HT (TPH2, red) cells in the DR projecting to the VTA (right panel, yellow; scale bar = 200 µm). Aq: cerebral aqueduct; IPR: interpeduncular nucleus. (C) Characterization of 5-HT_2C_R neurons (green) in the VTA shows dual-IF labelling (yellow, white arrows) with GABA using *vGat^Cre:mCherry^*(red) mice (left panel) or dopamine (tyrosine hydroxylase (TH), red; right panel; scale bars = 200 µm, magnified = 20 µm). (D) Most 5-HT_2C_Rs are co-expressed with GABA (*vGat^Cre:mCherry^*; n=4), not dopamine (TH, n=3) neurons in the VTA. (t(5) = 21.92, p <0.0001). (E) Model schematic of data related to Figure 5. (F) Whole-cell patch clamp electrophysiology in *dopamine transporter IRES-Cre*:*Rosa-tdTomato* mice (*Dat^Cre^:tdTom*) to visualize DA^VTA^ neurons (Scale bars: main image = 200 µm, magnified = 20 µm). **G-J**. 5-HT (10 µM) application increases spontaneous inhibitory postsynaptic currents (sIPSCs) onto DA^VTA^ neurons as illustrated in: (G) example traces; (H) time course of changes in sIPSC frequency (shaded bars, 5-HT application); (I) mean sIPSC frequency (t(6) = 3.335, p = 0.0157, n/N = 7/4); (J) sIPSC amplitude is not significantly different after 5-HT application (t(6) = 0.02402, p = 0.9816, n/N = 7/4). **K-N**. Application of 5-HT_2C_R antagonist RS 102221 (1 µM) prevented 5-HT (10 µM) induced changes of sIPSCs onto DA^VTA^ neurons as illustrated in: (K) example traces; (L) time course of sIPSC frequency changes (shaded bars, drug application); (M) mean sIPSC frequency (t(6) = 0.3260, p = 0.7555, n/N = 7/4); (N) sIPSC amplitude (t(6) = 1. 244, p = 0.2597, n/N = 7/4). **O-R**. Application of 5-HT_2C_R agonist WAY 161503 (30 µM) increases sIPSCs onto DA^VTA^ neurons as illustrated in: (O) example traces; (P) time course of sIPSC frequency (shaded bars, WAY 161503 application); (I) mean sIPSC frequency (t(7) = 4.628, P = 0.0024, n/N = 8/4); (R) sIPSC amplitude is not significantly different during WAY 161503 application (t(7) = 1.540, P = 0.1674, n/N = 8/4).

We therefore tested the effects of 5-HT_2C_R ligands on DA^VTA^ neurons using *dopamine transporter IRES-Cre* mice crossed with *Rosa-tdTomato* (*DAT^Cre:tdTom^*) mice to visualize DA^VTA^ neurons (**Figure S3A**). Evoked firing of tdTomato fluorescently identified DA^VTA^ neurons was recorded using whole-cell patch-clamp electrophysiology following bath applied 5-HT ligands. We examined whether 5-HT_2C_R antagonist RS 102221 could inhibit changes in firing produced by 5-HT (**Figure S3B-C**). Application of RS 102221 alone (102.5 ± 1.6%) did not alter firing rate of DA^VTA^ neurons, nor was this different after application of 5-HT (114.9 ± 9.3%). To test whether firing activity was altered by action at 5-HT_2C_Rs, the selective 5-HT_2C_R agonist WAY 161503 (30 μM) was bath applied, and this significantly decreased DA^VTA^ neuron firing (aCSF: 101.5 ± 0.7%, WAY: 63.2 ± 7.7%; **Figure S3D-F**). These findings suggest that activation of 5-HT_2C_Rs decreases firing activity of DA^VTA^ neurons.

Given that 5-HT_2C_Rs are primarily G_q_-coupled and highly expressed on GABA^VTA^ neurons, a reduction in DA^VTA^ neuron firing could be due to increased inhibitory input. We next tested how 5-HT alters spontaneous inhibitory postsynaptic currents (sIPSCs) within DA^VTA^ neurons (**Figure 5F**). Application of 5-HT increased sIPSC frequency onto DA^VTA^ neurons from 2.2 ± 0.4 Hz to 3.6 ± 0.6 Hz (**Figure 5G-I**). 5-HT did not alter sIPSC amplitude (aCSF: 24.5 ± 2.1 pA, 5-HT: 25.4 ± 3.4 pA, **Figure 5J**). To determine whether this effect was blocked by the 5-HT_2C_R antagonist, RS-102221 was bath applied with 5-HT to DA^VTA^ neurons. Application of RS-102221 prevented the increase in sIPSC frequency with 5-HT (aCSF: 1.9 ± 0.3 Hz, RS + 5-HT: 1.9 ± 0.3 Hz, **Figure 5K-M**). Again, there was no change in amplitude (aCSF: 34.7 ± 8.3 pA, RS + 5-HT: 33.1 ± 7.2 pA, **Figure 5N**). We then examined the effect of the selective 5-HT_2C_R agonist, WAY 161503 on sIPSCs to DA^VTA^ neurons. Application of WAY 161503 increased sIPSC frequency onto DA^VTA^ neurons from 2.8 ± 0.3 Hz to 4.1 ± 0.6 Hz (**Figure 5O-Q**) but did not alter sIPSC amplitude (aCSF: 33.9 ± 4.6 pA, WAY: 30.2 ± 2.5 pA, **Figure 5R**). Taken together, these results indicate that 5-HT released from DR neurons activates 5-HT_2C_R in GABA^VTA^ interneurons that in turn silence the activity of DA^VTA^ neurons.

### 5-HT2CRVTA neuronal activity modulates age-related PA

Following this ex vivo analysis, we established the effect of 5-HT_2C_R^VTA^ activation *in vivo* on PA levels. This was achieved by infusing AAV8-hSyn-DIO-hM3DGq (5-HT_2C_R^VTA^:hM3Dq mice) or control AAV8-hSyn-DIO-mCherry (5-HT_2C_R^VTA^:mCherry) into the VTA of *Htr2c-IRES-Cre* mice crossed with *Rosa-YFP* mice (*5-HT_2C_R^Cre:YFP^*) (**Figure 6A-B**). CNO (1 mg/kg, i.p.) treatment to young adult 5-HT_2C_R^VTA^:hM3Dq mice to activate 5-HT_2C_R^VTA^ neurons produced aging-related PA characteristics. This was apparent when mice were normally most active during the first 6 hours of the dark cycle, reducing home cage activity patterns visualized with heat maps (**Figure 6C**), number of beam breaks (**Figure 6D**), distance travelled (**Figure 6F**) and ambulation speed (**Figure 6G**) compared with vehicle administration. No differences in time spent inactive or food intake were observed (**Figure 6E, H**). These data are consistent with those obtained when activating 5-HT^DR^ neurons and illustrate that the exclusive activation of 5-HT_2C_R^VTA^ neurons promotes an age-related PA signature.

**Figure 6.**
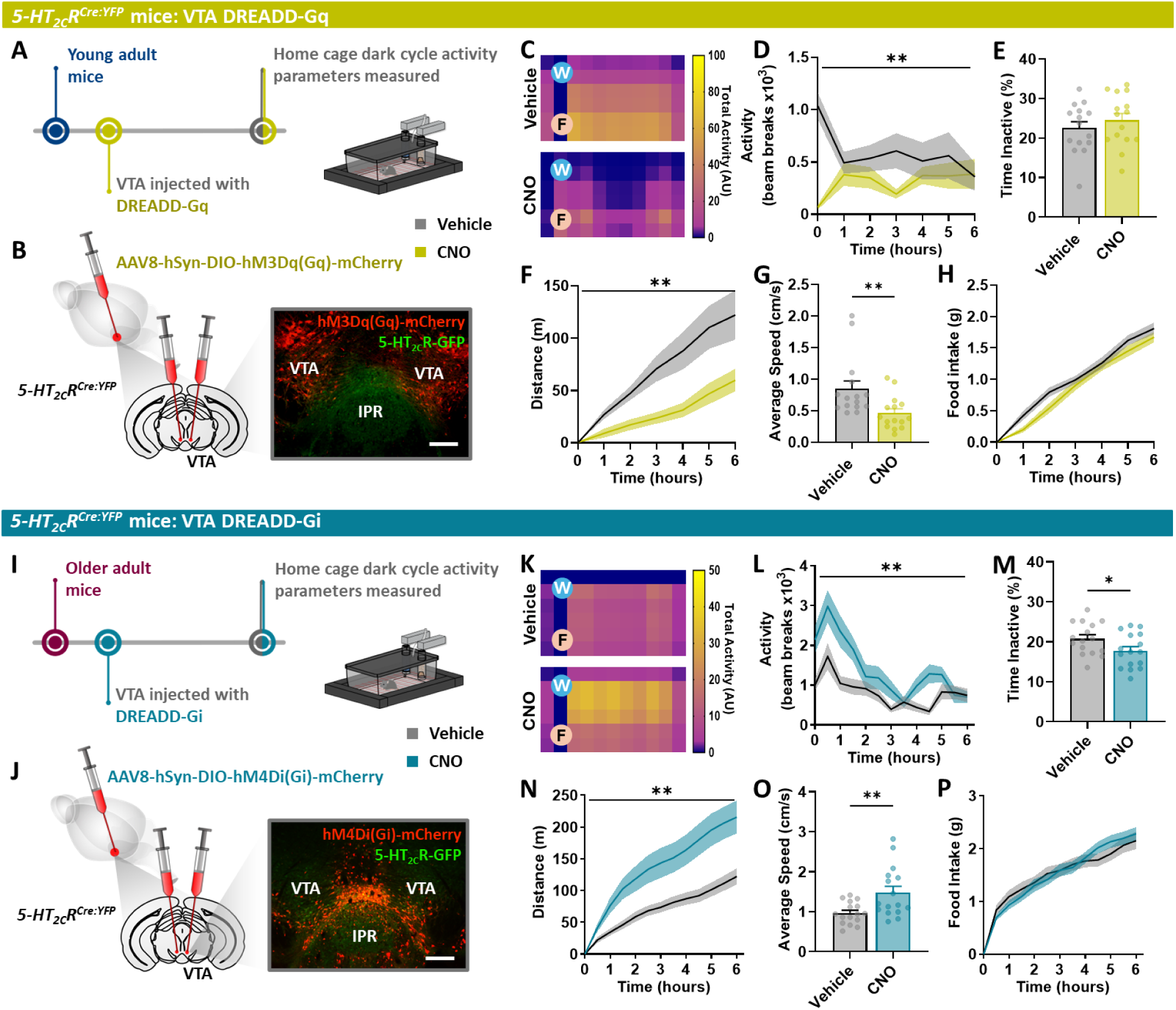
5-HT_2C_R^VTA^ neurons modulate age-related physical activity levels. **A-H.** (A) Schematic of chemogenetic study design in young adult *5-HT2CR^Cre:YFP^* mice (5 months old, n=15). (B) Schematic of VTA injection site and representative immunofluorescence (IF) image of AAV8-hSyn-DIO-hM3Dq(Gq)-mCherry (red) co-expressed (yellow) with 5-HT2CR-GFP neurons (green, scale bar = 200 µm). Treatment with designer drug clozapine-n-oxide (CNO; 1 mg/kg, i.p) produced an older adult profile in young adult mice compared to vehicle (saline) as illustrated by: (C) representative home cage ambulation heatmaps (W, water; F, food); (D) decreased total activity in beam breaks (RM ANOVA Treatment: F_(1,14)_ = 9.220, p = 0.0089); (E) no change in percentage time spent inactive (t(14) = 0.9370, p = 0.3646); (F) reduced cumulative distance (RM ANOVA Treatment: F_(1,14)_ = 13.93, p = 0.0022); (G) decreased average ambulatory speed (t(14) = 3.016, p = 0.0093); (H) no difference in food intake (RM ANOVA Treatment: F_(1,14)_ = 3.962, p = 0.0664). **I-P**. (I) Schematic of chemogenetic study design in older adult *5-HT2CR^Cre^* mice (12-16 months old, n=16). (J) Schematic of VTA injection site and representative immunofluorescence image of AAV8-hSyn-DIO-hM4D(Gi)-mCherry (red) co-expressed (yellow) with *5-HT2CR^Cre-YFP^*-GFP neurons (green, scale bar = 200 µm). Treatment with designer drug clozapine-n-oxide (CNO; 1 mg/kg, i.p) produced a younger adult profile in older adult mice compared to vehicle as illustrated by: (K) representative home cage ambulation heatmaps (W, water; F, food); (L) increased total activity beam breaks (RM ANOVA Treatment: F_(1,15)_ = 12.18, p = 0.0033); (M) reduced percentage time spent inactive (t(30) = 2.070, p = 0.0472); (N) increased cumulative distance (RM ANOVA Treatment: F_(1,15)_ = 11.01, p = 0.0047); (O) faster average ambulatory speed (t(15) = 3.285, p = 0.0050); (P) no difference in food intake (RM ANOVA Treatment: F_(1,15)_ = 0.0005, p = 0.9434).

To test the bidirectionality of this circuit, we injected AAV8-hSyn-DIO-hM4D(Gi)-mCherry (5-HT_2C_R^VTA^: hM4Di) or mCherry into the VTA of older adult mice to reversibly inhibit 5-HT_2C_R^VTA^ neuron activity (**Figure 6I-J**). CNO (1 mg/kg, i.p.) treatment in older adult 5-HT_2C_R^VTA^:hM3Di mice to dampen 5-HT_2C_R^VTA^ neuron activity promoted a more youthful PA profile. Specifically, CNO significantly increased dark cycle home cage PA as visualized in heat maps (**Figure 6K**) and quantified by beam breaks (**Figure 6L**) compared with vehicle administration. Further, CNO significantly reduced average time spent inactive (**Figure 6M**), increased average daily distance travelled (**Figure 6N**) and elevated ambulation speed (**Figure 6O**). CNO treatment did not influence food intake (**Figure 6P**). CNO did not alter behavior in control 5-HT_2C_R^VTA^:mCherry mice (**Figure S4**). These data illustrate that adjusting the tone of 5-HT_2C_R^VTA^ neuron activity produces a more youthful PA profile in older adult mice and an older adult PA pattern in younger adult mice, signifying that age-related PA characteristics are reversible and manipulable.

### Permanent 5-HT_2C_R^VTA^ neuronal silencing restores youthful activity characteristics in older adult mice

We extended these studies to examine the prolonged effect of preventing 5-HT^DR^ activity at 5-HT_2C_R^VTA^ neurons. *5-HT_2C_R^Cre:YFP^* mice were bilaterally injected in the VTA with an AAV5 expressing Cre-dependent flex-taCaspase-3-TEVp + AAV-mCherry (5-HT_2C_R^VTA^:Cas3) to selectively ablate 5-HT_2C_R^VTA^ expressing cells in older adult (12-18 months old) mice. ^60^ Age-matched control mice were treated with AAV-mCherry (5-HT_2C_R^VTA^:mCherry) (**Figure 7A-B**). Mice with ablated 5-HT_2C_R^VTA^ neurons appeared grossly normal behaviorally and ate normally (**Figure S5**). The most striking change observed in older adult 5-HT_2C_R^VTA^:Cas3 treated mice was a restoration of a more youthful PA profile, assessed both within the dark cycle (**Figure 7**) and as daily PA (**Figure S5**). Specifically, 5-HT_2C_R^VTA^:Cas3 mice engaged in more PA in the home cage as illustrated with heat maps (**Figure 7D**) and beam breaks (**Figure 7E**) compared to older adult control mice with functional 5-HT_2C_R^VTA^ neurons. Ablating 5-HT_2C_R^VTA^ neurons also reduced time spent inactive (**Figure 7F**), increased distance travelled (**Figure 7G**) and augmented ambulation speed (**Figure 7H**) compared control mice. This prolonged increase in home cage activity was associated with reduced body weight (**Figure 7J**) and body fat (**Figure 7K**) and improved lean mass in 5-HT_2C_R^VTA^:Cas3 treated mice compared to pre-treatment levels (**Figure 7L**). Given the increase in PA and changes in body composition, we measured maximal muscle strength to evaluate improvement in neuromuscular motor function.^61, 62^ Our results showed that ablation of 5-HT_2C_R^VTA^ neurons led to improved forelimb muscle strength compared to pretreatment, whereas non-ablated control mice showed no changes in strength. (**Figure 7M**). Post-mortem analysis confirmed significantly fewer 5-HT_2C_R^VTA^ neurons in 5-HT_2C_R^VTA^:Cas3 mice compared to control AAV-mCherry mice (**Figure 7C**). These data illustrate that preventing 5-HT signaling at 5-HT_2C_R^VTA^ neurons in older adult mice is sufficient to significantly reverse age-related activity/inactivity and associated changes in body weight, body fat, lean mass, speed and grip strength.

**Figure 7.**
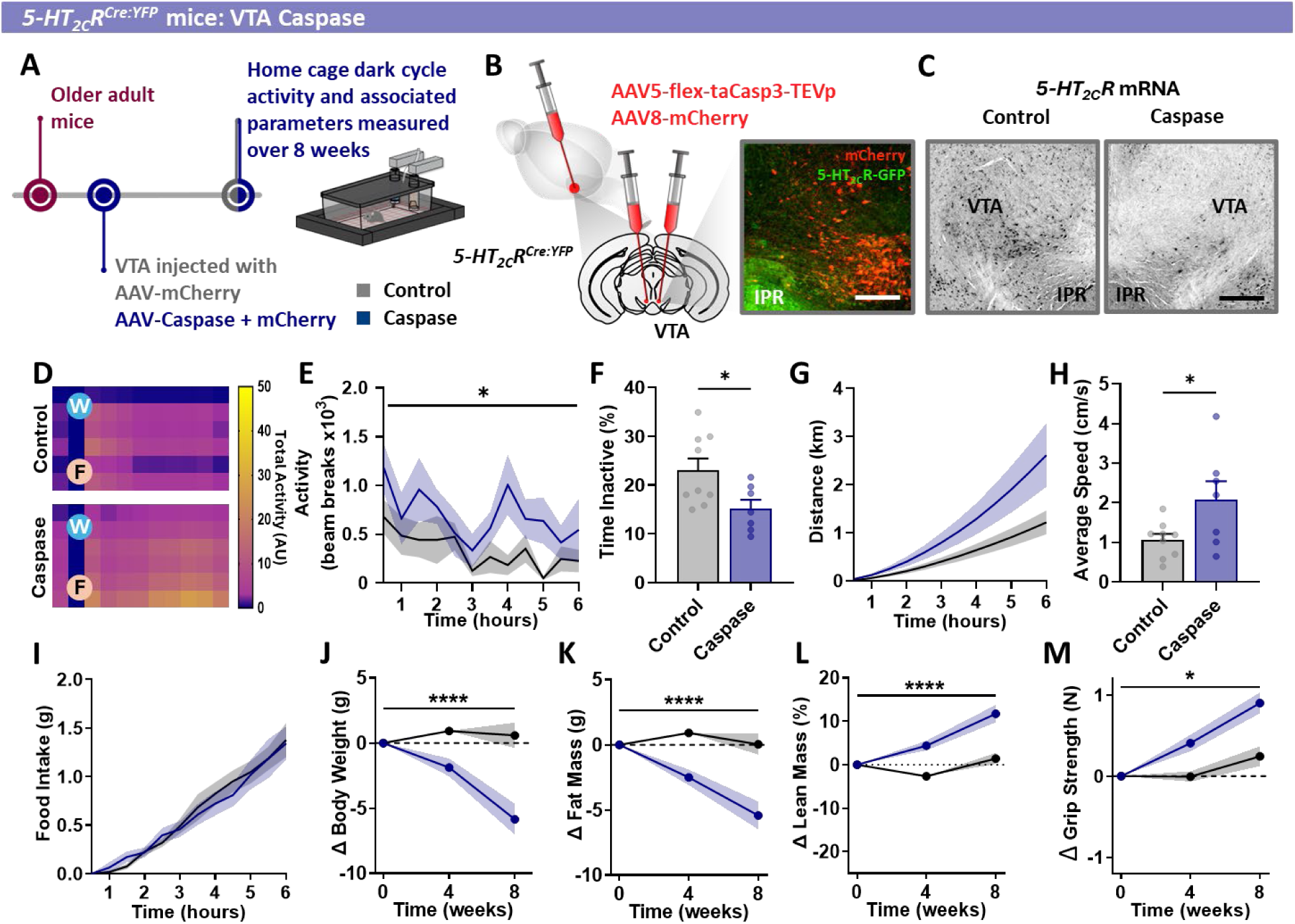
5-HT_2C_R^VTA^ neuron ablation restores a youthful activity profile in older adult mice. (B) Schematic of 5-HT2CR^VTA^ cell ablation study design in older adult *5-HT2CR^Cre:YFP^* mice (12-18 months old). (C) Schematic detailing co-injection of Caspase 3 and mCherry control into the VTA and representative immunofluorescence image of mCherry (red) with *5-HT2CR^Cre-YFP^*-GFP neurons (green, scale bar = 100 µm). (D) Representative 5-HT2CR^VTA^ ablation (right panel) as compared with controls (left panel) using *5-HT2CR* mRNA in situ hybridization histochemistry knockdown. Scale bar = 100 µm. **D-M**. Compared to control mice (n=9), permanently silencing 5-HT2CR^VTA^ neurons in older adult mice (n=7) restored younger adult physical activity phenotype as illustrated by: (E) representative home cage ambulation heatmaps (W, water; F, food); (F) increased home cage total activity measured in beam breaks (RM ANOVA Treatment; F_(1,11)_ = 8.665, p = 0.0134); (G) reduced percentage time spent inactive (t(14) = 2.437, p = 0.0287); (H) increased cumulative distance travelled (RM ANOVA Treatment: F_(1,14)_ = 4.262, p = 0.0580); (I) faster average ambulatory speed (t(14) = 2.325, p = 0.0356); (J) no difference in food intake (RM ANOVA Treatment: F_(1,11)_ = 0.02274, p = 0.829); (K) reduced body weight (Time*Treatment; F_(2,37)_ = 14.80, p < 0.0001); (L) decreased fat mass (ANOVA Time*Treatment: F_(2,37)_ = 15.82, p < 0.0001); (M) increased lean mass (ANOVA Time*Treatment: F_(2,37)_ = 18.35); increased grip strength (ANOVA Time*Treatment: F_(5,76)_ = 2.626, p = 0.0303).

## DISCUSSION

Physical inactivity progressively increases with age and is a major modifiable determinant of disease, disability, and mortality.^63, 64^ Here we characterized across species, older human adults (50–70 years) and life-phase–matched mice and found that they exhibited parallel reductions in daily distance and ambulatory speed alongside higher adiposity, signifying an intrinsic biological influence on midlife inactivity. Further, we identified a 5-HTergic mechanism that biologically programs the age-related decline in voluntary PA and demonstrate that targeted inhibition of this pathway promotes youthful activity and strength in older adult mice.

Although 5-HT^DR^ neurons arise from a shared developmental lineage,^65^ they are not homogenous; rather they comprise multiple physiologically and molecularly distinct subtypes.^41, 66-70^ Anatomical tracing studies indicate that different DR subregions preferentially target specific brain areas,^67, 68, 71^ and whole-brain input-mapping reveals corresponding diversity in presynaptic inputs to these 5-HT^DR^ neuron subgroups.^57, 66, 72^ Recent studies have employed viral-genetic tools to parse 5-HT^DR^ neurons according to their projection destinations, electrophysiological profiles, and roles in behavior.^70^ These approaches have begun to resolve longstanding contradictions and debate within the field.

While roughly 70% 5-HT^DR^ neurons are not glutamatergic,^73^ the subpopulation of dual 5-HT-glutamate (glutamate via vesicular glutamate transporter 3; VGLUT3) neurons projecting to the VTA predominantly establish asymmetric synapses on DA^VTA^ neurons.^74^ In contrast, non-glutamatergic 5-HT^DR^ projecting VTA neurons establish symmetric synapses primarily on VTA non-DA neurons (presumably GABAergic).^74^ GABA^VTA^ neurons provide a local inhibitory input to DA^VTA^ neurons.^75, 76^ Optogenetic activation of VGLUT3^DR^ neurons increases locomotor activity^77^ and in turn, activation of DA^VTA^ neurons stimulates PA.^52^ Here we show that chemogenetic activation of 5-HT^DR^ neurons inhibits home cage physical activity. This dual DR 5-HT-glutamatergic and 5-HT-only → VTA connectivity enables 5-HT^DR^ to modulate locomotion and motivational drive through both direct excitatory and indirect inhibitory mechanisms onto DA neurons. Our results suggest that, during aging, enhanced 5-HTergic tone during wakefulness biases this circuit toward inhibition through 5-HT_2C_R-mediated activation of GABA^VTA^ neurons, thereby damping DAergic drive and voluntary PA.

This interpretation is consistent with our findings illustrating that chemogenetic activation of 5-HT^DR^ neurons in young adult mice reproduced an older adult PA signature, whereas blocking 5-HT action at 5-HT_2C_Rs prevented age-related declines in PA. Immunohistochemical mapping and *ex vivo* electrophysiology revealed that 5-HT_2C_Rs are predominantly expressed on VTA^GABA^ neurons rather than DA^VTA^ neurons. Further, we found that 5-HT_2C_Rs indirectly inhibit DA^VTA^ neurons via activation of GABAergic interneurons. Bidirectional manipulation of 5-HT_2C_R^VTA^ neurons revealed a behavioral effect of this mechanism *in vivo*: 5-HT_2C_R^VTA^ neuron activation suppressed home cage locomotion in young adults, while inhibition or ablation restored youthful distance, speed, and grip strength in older adult mice. This finding explains pharmacological evidence that 5-HT_2C_Rs signaling constrains locomotion, whereas antagonists or gene knockout increase DA signaling and locomotor output^45-47, 78-81^ by discovering a causal, circuit-level specificity to 5-HT^DR^ → 5-HT_2C_R^VTA^ gating.

Aging of the 5-HT system likely contributes to this shift in locomotor and speed set-point. Further illustrating the cross-species consistency of this circuit, as we report in mice, no difference in the number of 5-HT^DR^ neurons have been observed in older versus younger people.^82^ We report a phase-dependent remodeling of 5-HT^DR^ neuron firing with age—elevated during the active/wakeful period—consistent with the inverse relationship between 5-HT tone and locomotion.^37^ Our data suggest that the activity of the 5-HTergic system changes with aging, and the augmented inhibitory drive via 5-HT_2C_Rs slows PA. Targeting this progressively maladaptive signaling may thus restore motivational tone and physical mobility.

This study focused on naturalistic, spontaneous PA, which carries independent prognostic value beyond organized MVPA. Specifically, walking is the most prevalent form of daily PA in adults across populations,^83^ and speed and cadence are strong predictors of longevity, cardiovascular health, and cognitive resilience.^32, 84, 85^ Importantly, improvements in everyday movement—even without structured exercise—are associated with substantial reductions in mortality risk. Thus, uncovering a neurobiological pathway that governs spontaneous PA is highly relevant to public health, as it may enable restoration of PA patterns in midlife.

Collectively, these findings fill an essential gap by defining a brain circuit that programs the age-related decline in voluntary PA and highlighting 5-HT_2C_R^VTA^ neurons as a tractable node to restore mobility in midlife. The development of VTA targeted pharmacological 5-HT_2C_Rs antagonists could provide a therapeutic strategy to enhance voluntary movement and counter conditions linked to sedentary aging such as sarcopenia, metabolic dysfunction, and cognitive decline. Because our manipulation also improved muscle strength and body composition, it suggests that central 5-HTergic tone directly influences not only movement motivation but also physical capability.

Several limitations should be acknowledged. First, the human study was cross-sectional and conducted within a single cultural and geographic population. Our data were in keeping, however, with other, larger multinational and multiethnic studies demonstrating the trajectory of midlife PA decline.^86, 87^ Second, caloric intake was self-reported, and recall bias cannot be excluded, although objective accelerometer data strengthen our conclusions about PA patterns. Chemogenetic and pharmacological interventions, while insightful, may influence off-target circuits. Our electrophysiological recordings were performed in brain slices, which remove modulatory inputs that may influence *in vivo* firing patterns.

In summary, this study defines a brain circuit that governs the natural decline in PA with age. By pinpointing 5-HTergic 5-HT_2C_R signaling in the VTA as a pivotal and modifiable brake on spontaneous PA, our findings reveal that the drive to be physically active is not merely a lifestyle choice but a biologically regulated state that can be reactivated. Understanding and targeting this mechanism provides a new avenue to preserve mobility, independence, and quality of life in an aging population – transforming the biology of inactivity into an opportunity for healthy longevity.

## METHODS

### Subjects

#### Study participants

As part of The Full4Health project,^53^ 72 healthy male and female participants within two age groups – young (17-30 years old, n=44) and older (50-70 years old, n=28) adults were recruited in Scotland by public advertisement on the radio, newspaper and social media. To form this healthy, non-athletic cohort, subjects without the following exclusion criteria were enrolled: smokers; morbid obesity (BMI>40 kg/m^2^); pregnancy; obesity of known endocrine origin; neurological disorders; medication known to influence appetite; self-reported fever/systemic infection; participation in medical or surgical weight loss program within 1 month of selection; history of cerebrovascular disease; current major depressive disorder; history of cardiovascular disease; chronic obstructive pulmonary disease; an allergy to any of the test drink components of The Full4Health project and partaking in > 6 hours of vigorous PA per week. This study was performed in accordance with the Declaration of Helsinki.^88^ Ethical approval in Aberdeen was granted by the National Health Service North of Scotland Research Ethics Service. All participants provided written informed consent before entering the study. Data from 17–18-year-olds were collected at schools if individuals were still in school with appropriate safeguarding, whereas adults attended the Rowett Institute, University of Aberdeen, Scotland for assessment.

#### Animal models and housing conditions

Young (1-6 months old) and older (12-24 months old) adult male and female mice on a C57BL/6 background were used as indicated in figure legends. This included wild type, *Pet^YFP^*,^89^ Tph2-iCreER (#016584, JAX), DAT-Cre (#006660, JAX)^90^, tdTomato-LoxP (Ai9) (#007909, JAX)^91^, Vgat-Cre (#016962, JAX),^92^ and *5-HT_2C_R^Cre:YFP^* ^93, 94^ mice. Mice were group-housed with *ad libitum* access to food and water in a 12-hour light/12-hour dark cycle in a temperature-controlled environment (20-22°C). Mice were maintained on a chow diet (CRM(P) 801722, Special Diet Services in Aberdeen and diet 5062 Pico-Vac from LabDiet). All procedures were carried out in accordance with the U.K. Animals (Scientific Procedures) Act 1986 or Canadian Council for Animal Care and were approved by local university ethical approval at University of Aberdeen or University of Calgary, respectively.

### Body measurements

#### Participants

Height, body weight and body composition were measured in the fasted state and after voiding as described previously.^95^ Height was measured to the nearest 0.1 cm using a portable stadiometer (Model 213, SECA, Germany).^96^ Body weight, measured to the nearest 0.1 kg, and body composition were assessed using a multi frequency segmental body composition analyzer (Model BC-418-MA, Tanita, Japan). Body mass index (BMI) was calculated for each participant and compared against the age and gender-matched thresholds for normal-weight and overweight, as defined by the World Health Organization^97^. In addition, visceral fat and trunk fat percentages were measured in the supine position using bioelectrical impedance analysis technology (AB 140 Viscan, Tanita, Japan).

#### Mice

Body weight and composition in mice was measured in a fed state. Body weight was measured to the nearest 0.01g using animal weighing scales (Ohaus Adventurer Pro, USA). Body composition, including fat mass and fat-free mass, was measured to the nearest 0.01g by EchoMRI-500 Body Composition Analyzer (Zinsser Analytic GmbH, UK) and an average of three repeat scans was taken.

### PA and food intake

#### Participants

Free-living PA was assessed over seven consecutive days by Actigraph GT3X+ accelerometers (Actigraph, LLC, Fort Walton Beach, Florida) worn on the hip. Participants were shown how to wear the accelerometer and left the research site wearing it in the correct position. Formal data collection began the next day. Upon completion, participants returned the devices to the research team. Bodily movements were measured on three axes with a frequency of 30Hz and epoch length of one minute. Cadence was classified into categories of 0-29, 30-59, 60-89, 90-119 and >120 steps per minute. The proportion of time in each cadence category was calculated for each participant, with an assumption that activity was in the lowest category when the device was switched off. Distance covered (km) was estimated as 0.43 x height(m) x steps. Walking speed was estimated from mean distance per minute in all intervals of 5 or more minutes in which the cadence was at least 60 steps per minute and converted to km/hour. An assessment of habitual food intake was obtained using 24-hour dietary recall during each of the four experimental visits to the Rowett Institute or school.

#### Mice

Locomotor activity and food intake measurements were performed using TSE PhenoMaster System cages (TSE, Germany). Mice were habituated to the cages for at least 5 days before the experiment commenced. For studies involving administration of drugs, food was removed 30 minutes prior to the onset of the dark cycle and mice were injected with vehicle or drug. Locomotor activity, mouse movement, speed and home cage location were retrieved from time-stamped beam breaks from 21 lasers evenly distributed around the cage (8 in cage width (x-axis) and 13 in cage length (y-axis)). Total locomotor activity was calculated as summatory of beam breaks per unit of time; distance travelled was retrieved directly from the TSE PhenoMaster System measurement combining beam breaks and distance algorithm; speed was calculated as an average speed from 1 minute interval speed data, measured in cm/s, collected directly from the TSE PhenoMaster System; inactivity was calculated from 1 minute interval activity sensor data as a percentage of time when non-ambulatory activity was detected. Activity heatmaps were generated using a grid depicting infrared activity sensor beam breaks, multiplied for each square, and normalized to the maximum beam break level. Beam break crossings were computed over a period of 6 hours and normalized to allow comparison. Activity heatmaps were generated using GraphPad Prism. Food intake was measured directly from continuous food sensor measurements from the TSE PhenoMaster System and plotted as cumulative food intake.

### Grip strength test

A grip strength test was used to measure the neuromuscular function of mice. Briefly, mice were tail suspended and allowed to grip a horizontal metal grid with their forelimbs. A load cell sensor attached to the grid and controlled using Arduino program detected the maximal pulling force applied before the mouse released the grid. Each mouse was tested 3 times in a single daily session, once a week over 8 weeks. Maximal pulling force for each trial was averaged for each animal and session.

### Geometric graphing

Geometric graphing was used to identify similarities in the aging metabolic signature between human participants and mice (Figure 1 M-N). Each subject was considered as a point in 6-dimensional space, with each dimension a particular property of the subject. The values of the properties were normalized as percentages relative to the average of each property, so for example, “distance” in all would be on the same scale for participants and mice. The 6-dimensional values created data “sphere,” and a threshold distance of 24 between spheres was chosen (by trial and error) to best reflect the structure of the data. When two spheres intersect, the two subjects representing these spheres are connected. To determine intersections of spheres, the standard Euclidean distance was used. Such a collection of objects and connections among the objects describes a graph, a common tool^98, 99^ of topological data analysis and of biology.^100^ The graph was visualized using Mathematica (Wolfram Research Inc., 2020), with the SpringElectricalEmbedding graph layout. This layout minimizes total energy, considering connections as springs and subjects as charges. Finally, the graph was colored to help visually compare the observed data with the topological separation of the branches of the graph.

### Viral vectors and stereotaxic surgery

Cre-dependent DREADD viral vectors include adeno-associated virus (AAV) AAV8-hSyn-DIO-hM3D(Gq)-mCherry (1.83x1012 gc/ml), AAV8-hSyn-DIO-hM4D(Gi)-mCherry (1.7x1012 gc/ml), and AAV8-hSyn-DIO-mCherry (3.6x1012 gc/ml) were a generous gift from Prof Bryan Roth (Addgene plasmid # 44361, # 50459, # 14472).^101^ Cre-dependent AAV5-Flex-taCasp3-Tevp (Vector Core, AV5760)^60^ was used to program cell death, co-infused (10:1) with pAAV-hSyn-mCherry (AAV8) (Addgene, 14472) to visualize injection site. Retrogradely transported canine-adenovirus (CAV)-2-green-fluorescent protein (GFP) was used to track 5-HT cells projecting to the VTA (Universitat Autonoma # CBATEG-493).

For the delivery of viral particles into the discrete brain regions, stereotaxic surgery was adapted from previous studies.^102^ Briefly, mice were anaesthetized with isoflurane, the top of the head was shaved and they were placed in a stereotaxic frame (David Kopf Instruments, CA, USA). A longitudinal incision was made in the skin to expose the surface of the skull and hole was drilled at the specific coordinates from bregma (anteroposterior, dorsoventral and lateral to midline in mm) for the VTA (-3.16; -4.5; ±0.56) and for the DR:4.7; -3.5; 0). Using a pulled glass capillary (40μm tip diameter) (G1, Narishige, UK) and a pneumatic microinjector (IM-11-2, Narishige, UK), 500 nl of viral preparation was bilaterally injected into the VTA or into the DR at a flow rate of 50nl/minute. The capillary was left in the injection site for 5 minutes to allow diffusion before being slowly removed to avoid dispersion to neighboring regions. Injection validation was performed with immunohistochemistry (IHC). Mice with missed injections were excluded from analyses.

### Electrophysiology

Mice were deeply anaesthetized, decapitated, and horizontal midbrain sections (250 μm) containing the DR or VTA were cut using a vibratome (VT1200, Leica Microsystems, Nussloch, Germany). Slices were recovered in warm NMDG solution (32 °C) saturated with 95% O_2_–5% CO_2_ for 10 min before being transferred to a holding chamber containing artificial cerebrospinal fluid (ACSF) and equilibrated with 95% O_2_/5% CO_2_ for at least 45 min before recording. Slices were transferred to a recording chamber on an upright microscope (Olympus BX51WI, Evident Scientific, Quebec, Canada) and continuously superfused with ACSF (2 ml/min, 34 °C).

*ePET^YFP^* mice (n = 8) were used to establish the intrinsic activity 5-HT^DR^ neurons with age. To achieve this, a single-cell *ex vivo* electrophysiology study in younger (2-4 months) and older (12-24 months) male and female mice was performed using methods reported earlier.^103^ The animals were culled by cervical dislocation followed by decapitation, and coronal brain sections (180 µm) containing the dorsal raphe were cut in ice-cold ‘slicing’ solution (in mM: 2.5 KCl, 1.3 NaH_2_PO_4_, 26 NaHCO_3_, 213.3 sucrose, 10 glucose, 2 MgCl_2_, 2 CaCl_2_, bubbled with 5%CO_2_/95%O_2_) using a vibratome (Campden Instruments 7000smz-2). The brain sections were incubated at 36 °C in artificial cerebrospinal fluid (ACSF; in mM: 125 NaCl, 2.5 KCl, 1.2 NaH_2_PO_4_, 21 NaHCO_3_, 1 glucose, 2 MgCl_2_, 2 CaCl_2_) for at least 45 minutes before being transferred to a holding chamber containing ACSF at room temperature and bubbled with 5%CO_2_/95%O_2_. Slices were transferred to a recording chamber on an upright microscope (Olympus BX51WI) and continuously superfused with ACSF at room temperature. Dorsal raphe 5-HT (5-HT^DR^) neurons were identified by yellow-green fluorescence. Recording pipettes (3–6 MΩ) were filled with a 150 mM NaCl solution. Firing activity was recorded with a MultiClamp 700B amplifier (Axon Instruments, Molecular Devices) using the cell-attached technique in voltage-clamp mode. To ensure that firing activity remained undisturbed the command potential was set to the value that gave a holding current of 0, and 3-minute-long traces were acquired.^104^ To define 5-HT^DR^ neurons activity during the dark cycle, mice were housed under a shifted light schedule (lights on at 11:45 pm, lights off at 11:45 am).

*DAT^Cre:tdTom^* mice (n=20) were used for pharmacology studies to define the effect of 5-HT_2C_R agonists and antagonists on DA^VTA^ neuron activity. Single-cell *ex vivo* electrophysiology was conducted using male and female mice (1–2 months old) using methods previously published.^105^ DA^VTA^ neurons were identified by red fluorescence and morphological and electrophysiological characteristics [fusiform shape, capacitance >50 pF, presence of H-current (Ih)]. Recording electrodes (3–5 MΩ) were filled with potassium-d-gluconate internal solution, (in mM): 136 potassium-d-gluconate, 4 MgCl_2_, 1.1 HEPES, 5, EGTA, 10 sodium creatine phosphate, 3.4 Mg-ATP, and 0.1 Na_2_GTP. Because most dopamine neurons ceased firing within 5 minutes of recording, current-step induced firing was used for all experiments. Firing activity was recorded in the current clamp mode. For current-step experiments, the membrane potential for each neuron was set to -60 mV by DC injection via the patch amplifier and a series of 5 current pulses (250 ms in duration, 5-25 pA apart, adjusted for each cell) were applied every 45 seconds. From this series of current steps, we then selected a current step that yielded 3-4 action potentials during the baseline period and used that step for the analysis. During the baseline period (first 10 minutes of the recording), all data points were normalized to the mean response averaged over 10 minutes. 3 neurons were excluded due to unstable baselines. Drug effects were calculated by comparing the response during the baseline/pre-drug application period to the response to this current step following drug application. For voltage clamp experiments, recording electrodes were filled with a cesium chloride solution (CsCl) internal solution consisting of (in mM) 140 CsCl, 10 HEPES, 0.2 EGTA, 1 MgCl_2_, 2 MgATP, 0.3 NaGTP, 5 QX-314-Cl. The junction potential of +4 mV for CsCl internal or +16.2 mV for KGluconate internal solution was not corrected for. Series resistance (6-20 MΩ) and input resistance were monitored on-line with a 5-mV depolarizing step (50 ms). Recordings exhibiting a >20% change in series resistance were discarded. GABA_A_ sIPSCs were selected for amplitude (>12 pA), rise time (<4 ms), and decay time (<10 ms). Experiments were performed during the animal’s light cycle.

### Immunohistochemistry (IHC)

Brain tissue was collected, prepared and processed for IHC as previously described.^58, 102, 106, 107^ In brief, mice were injected with a terminal dose of pentobarbital and perfused with phosphate buffered saline (PBS) (Gibco (Fisher), Cat#18912-014) followed by 50 ml of neutral buffered formalin 10% (Sigma, Cat# HT501320-9.5L). Brains were post-fixed in formalin for 24 hours and emersed in 30% sucrose (Sigma, Cat# D9663) for 24-48 hours. Tissue was sectioned at 25 μm on a freezing sliding microtome (Bright Instruments) and collected in 5 equal series. Fluorescent IHC was performed as followed: tissue was washed in PBS with 0.2% Tween (Sigma, Cat# P1379 and P9416), blocked using 5% normal donkey serum (Sigma, Cat# D9663) and 0.25% triton (Triton X-100, Fluka), and then incubated overnight with primary antibody in 2.5% normal donkey serum and 0.25% triton. Primary antibodies included: rabbit anti-RFP (1:2000, Rockland, Cat# 600-401-379, RRID:AB_2209751), chicken anti-GFP (1:1000, Abcam, Cat# ab13970, RRID:AB_300798) and goat anti-Tph2 (1:2,000, Abcam, Cat# ab121020, RRID:AB_10975512), goat anti-mCherry (1:1000, Sicgen, Cat# AB0040, RRID: AB_2333093), mouse anti-SR-2C(1:300, Santa Cruz, Cat# sc-17797, RRID:AB_628241), rabbit anti-TH(1:1000, Sigma, Cat# AB152, RRID:AB_390204).

Tissue was next washed in PBS with 0.2% Tween and incubated with the appropriate secondary antibody (Alexa Flour Donkey anti-rabbit 568 (1/500, ThermoFisher, Cat# A10042, RRID:AB_2534017), Alexa Flour Donkey anti-chicken 488 (1/500, Jackson ImmunoResearch, Cat# 703-545-155, RRID:AB_2340375), Alexa Flour Donkey anti-mouse 488 (1/500, Invitrogen, Cat# A21202, RRID: RRID:AB_141607), Alexa Flour Donkey anti-goat 488 (1/500, ThermoFisher, Cat# A11055, RRID:AB_2534102) or Alexa Fluor Donkey anti-goat 568 (1/500, ThermoFisher, Cat# A11057, RRID:AB_2534104) for 1 hour. DAPI (1:5,000) was with the secondary antibody for nuclear staining. Tissue was finally washed using PBS and 0.2% tween and mounted on a microscope slide (Epredia). Chromogenic IHC used primary goat anti-Tph2 (1/2000, Abcam, Cat# ab121020, RRID:AB_10975512) and rabbit anti-cFos (1/5000, Millipore, Cat# ABE457, RRID:AB_2631318) antibodies, secondary antibodies biotin-SP donkey anti-goat (1/500, Jackson ImmunoResearch, Cat# 705-065-147, RRID:AB_2340397) or Biotin-SP donkey anti-rabbit (1/500, Jackson ImmunoResearch, Cat# 711-065-152, RRID:AB_2340593), avidin-biotin complex (ABC) kit (Vector Laboratories, Cat# PK-6100, RRID:AB_2336819) and 3,3’-diaminobenzidine (DAB) kit (Vector Laboratories, Cat# SK-4100, RRID:AB_2336382). Images were acquired with Axioskop II microscope (Carl Zeiss, Germany) and Axio Vision Software (Carl Zeiss, Germany). Images were analyzed using ImageJ.^108^ DR and VTA were assigned according to Paxinos & Franklin mouse brain atlas and anatomical landmarks.^109^ mCherry or GFP expression was examined to identify injection site. Tph2 cell expression was determined as total number of cells expression Tph2 at bregma -4.60. 3D reconstruction of 5-HT^DR^ neurons was performed using Free-D software.^110^

### In situ hybridization histochemistry (ISH)

To validate the ablation of 5-HT_2C_R^VTA^ neurons in old mice we performed a post-mortem analysis of *5-HT_2C_R* mRNA expression pattern in the VTA using mRNA *in situ* hybridization adapted from previous works.^111^ Briefly, brains were perfused with RNase-free PBS-DEPC solution and fixed using sterile 4% paraformaldehyde, PFA. Brains where then dissected, post-fixed with PFA and cryoprotected with RNase-free 20% Sucrose solution. Brains were sectioned at 25μm, mounted in glass slides and kept at - 20C until. For mRNA hybridization, sections were cleaned in PBS-DPC and pretreated with consecutive solutions containing 1% NaHB4, 0.1M TEA and 0.25% acetic anhydride. Sections where then treated with SSC and incubated with pre-hybridization buffer. Sections where then incubated with DIG-labelled PCR-generated RNA probes against *5-HT_2C_R* mRNA (NM_008312.2) using validated primer pairs from Allen Brain Atlas (FWD: AGACCAGAATGATGCACATGAC and RV: GAGAGCTTGGAAGGATCTGAAA). Unbound probe was removed using stringency washes. Amplification of the signal was performed using TSA-Biotin kit and following manufacturer instructions. Visualization of the signal was performed using chromogenic staining with DAB as described above.

### Drugs

D-Fenfluramine (3 mg/kg; Tocris Bioscience, Cat# 2695; CAS: 3239-45-0) and clozapine-N-oxide (CNO) (1 mg/kg; Tocris Bioscience) were dissolved in vehicle (0.9% NaCl for infusion) and injected intraperitoneally 30 min before behavioral assays. Intraperitoneal injections were made of a volume of 10 ml/kg body weight. Tamoxifen (4-Hydroxy-tamoxifen dissolved in sunflower oil) (4 mg/kg, i.p., Sigma) was administered days 3, 7 and 11 post-stereotaxic surgery. *Ex vivo* applied drugs 5-HT (10 µM, Tocris Cat #3547), RS-102221 (1 µM, Tocris, Cat #1050), and WAY-161503 (30 µM, Tocris, Cat # 1801), were dissolved in DMSO (concentration 0.01%, Millipore Sigma, Cat #472301) and diluted to their final concentration in ACSF and bath applied to slices.

### Statistical analysis

Raw data were inputted into or transferred to Excel (Microsoft) files. Firing data and sIPSCs were analyzed with the MiniAnalysis program (Synaptosoft). Plots and statistical analyses were generated in GraphPad Prism 9.4 (GraphPad Software). The normality of data distribution was checked by Shapiro-Wilk W-test. For between study designed experiments the data was analyzed by parametric (Welch corrected or in case of equal variances Student t) or nonparametric (Mann-Whitney U) tests. For within-study designed experiments, paired t-test, Wilcoxon matched-pairs signed-rank test or repeated measures ANOVA for two-group comparisons were used. Activity levels were analysed by factorial ANOVA with terms for age, sex and their interaction. Where appropriate, data are presented as individual data points, and summary data are presented as mean ± standard error of the mean (SEM). Details of statistical tests results and number of animals can be found in the figure legends. Statistical significance was defined as *p <0.05, **p<0.01, ***p<0.001 and ****p<0.0001.

## ACKNOWLEDGEMENTS

The authors thank Dr. Lourdes Valencia Torres, Dr. Cristian Olarte Sanchez, Dr. Celine Cansell and staff within the University of Aberdeen Medical Research Facility and the Microscopy Facility staff for their technical assistance. The authors also wish to thank the Full4Health study research participants for taking part in the study. *Pet^YFP^*mice were generously provided by Prof Evan Deneris, Case Western Reserve University. Work was supported by the Biotechnology and Biological Sciences Research Council (BB/R01857X/1, BB/V016849/1 to LKH), University of Aberdeen Wellcome Trust Institutional Strategic Support Fund award (to L.K.H., S.L.B., P.B.M.d.M. and G.B.; 204815/Z/16/Z to L.K.H. and R.L.), The Royal Society of Edinburgh Research Reboot: COVID-19 Impact (#1122 to P.B.M.d.M.), Canada Research Chair Tier 1 (950-232211 to S.L.B.), Alberta Children’s Hospital Research Institute Postdoctoral Scholars Award (to DMN), and Scottish Government, Rural and Environmental Science & Analytical Services (RESAS) division (to G.H., A.M.J.). The Full4Health project was funded by the European Union’s Seventh Framework Programme FP7-KBBE-2010-4 under grant agreement No:266408 (to A.M.J.).

## CRediT AUTHOR CONTRIBUTIONS

**Conceptualization**: L.K.H and P.B.M.d.M.

**Methodology**: L.K.H., P.B.M.d.M., R.L., J.L., J.A.G., S.L.B.

**Software**: P.B.M.d.M, G.H., J.L., R.L., A.L-P.

**Validation**: P.B.M.d.M., G.S.B., J.A.G., S.L.B.

**Formal analysis**: L.K.H., P.B.M.d.M., A.L-P., S.M., Y.M., J.A.G., A.M., D.M.N., S.L.B.

**Investigation**: P.B.M.d.M., A.L-P., S.M., Y.M., A.M., D.M.N., J.A.G., J.L., C.C., M.A., G.S.B., A.L., G.H.

**Resources**: L.K.H, P.B.M.d.M., A.J.M., D.R.C., C.L.F., W.B., D.P., E.D.

**Data curation**. L.K.H., G.H., P.B.M.d.M., A.L-P, A.J.M, D.R.C., C.L.F., W.B., D.P.

**Writing-original draft**: L.K.H, P.B.M.d.M., A.L-P.

**Writing -Review & editing**: S.M., Y.M., A.M., D.M.N., J.A.G., J.L., C.C., D.R.C., C.L.F., W.B., D.P., M.A., G.S.B., A.L., E.D., G.H., R.L., A.M.J., S.L.B.

**Visualization**: A.L-P., P.B.M.d.M., A.M., D.M.N., J.A.G., J.L., G.H.

**Supervision**: L.K.H.

**Project Administration**: L.K.H.

**Funding acquisition**: L.K.H., P.B.M.d.M., G.B., A.J.M, S.L.B.

