## Supplemental Figures & Methods for "Brain serotonin circuit reverses decline in physical activity with age"

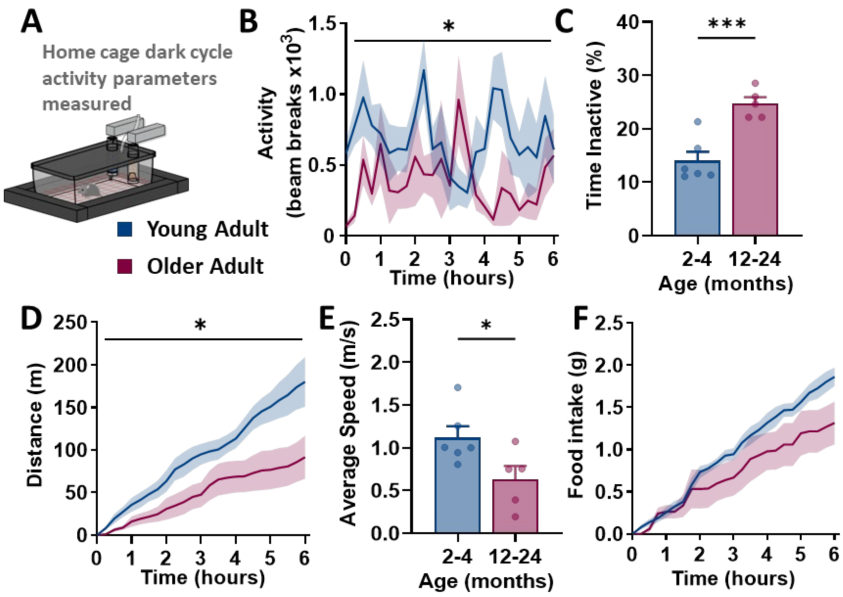


**Figure S1 pertaining to Figure 1 and 2. Decline in physical activity characteristics in older adult mice.**

(A) Schematic detailing study design and assessment of young adult (n=6) and older adult (n=5) C57BL/6 mice.

(B) Total activity in beam breaks in young adult and older adult mice (RM ANOVA Age: $F_{(1,9)} = 6.919$, $p = 0.0273$).

(C) Time spent inactive in young adult and older adult mice ( $t(9) = 4.996$, $p = 0.0007$).

(D) Cumulative distance travelled in young adult and older adult mice (RM ANOVA Age: $F_{(1,9)} = 5.681$, $p = 0.0410$).

(E) Average speed in young adult and older adult mice ($t(9) = 2.421$, $p = 0.0385$).

(F) Cumulative food intake in young adult and older adult mice (RM ANOVA Age: $F_{(1,9)} = 2.313$, $p = 0.1626$).

*p<0.05, **p<0.01, ***p<0.001.

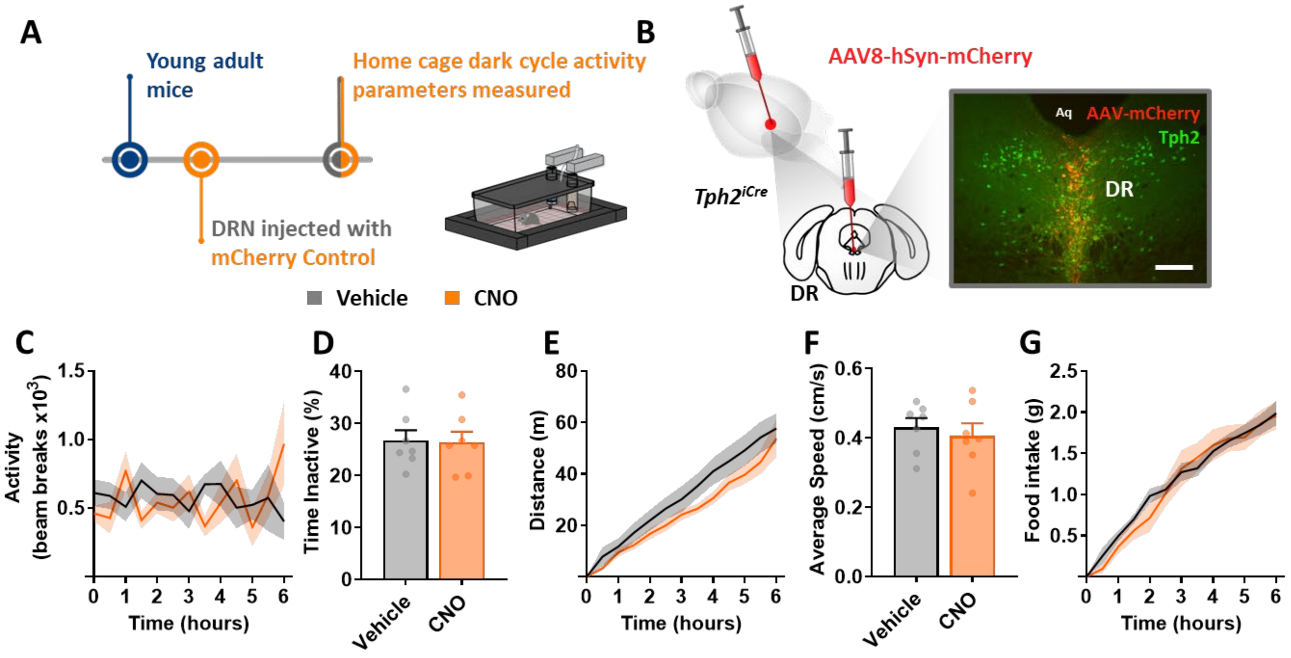


**Figure S2 pertaining to Figure 3. Chemogenetic *Tph2*[iCre] dorsal raphe (DR) control mice show no changes in physical activity.**

(A) Schematic detailing study design, stereotaxic injection of AAV-hSyn-mCherry control into the DR (5-HT$^{DR}$:mCherry) in young adult mice (n=7), and assessment with designer drug clozapine-n-oxide (CNO, 1 mg/kg, i.p.) treatment compared with vehicle (saline).

(B) Schematic detailing AAV-hSyn-mCherry injection site and immunohistochemistry (IF) image showing injection site (red) and Tph2-IF (green) expression and co-expression (yellow) in the DR. Scale bar = 100 µm. Cerebral aqueduct, Aq. No treatment differences were found including:

(C) total activity in beam breaks (RM ANOVA Treatment: $F_{(1,6)} = 0.04304$, $p = 0.8425$);

(D) time spent inactive ($t(6) = 0.09621$, $p = 0.9265$);

(E) cumulative distance (RM ANOVA Treatment: $F_{(1,6)} = 1.014$, $p = 0.3529$);

(F) average speed ($t(6) = 0.5134$, $p = 0.6260$);

(G) cumulative food intake of (RM ANOVA Treatment: $F_{(1,6)} = 0.09940$, $p = 0.7632$).

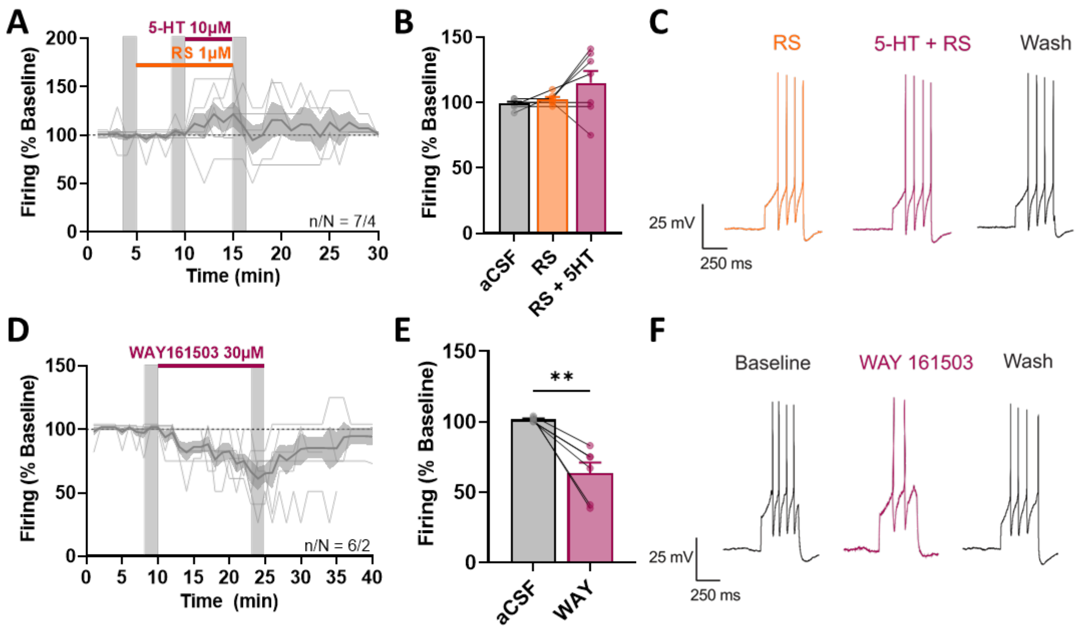


**Figure S3 pertaining to Figure 5. 5-HT$_{2C}$R$^{VTA}$ activation decreased the firing rate of DA$^{VTA}$ neurons.**

(A) Application of a selective 5-HT$_{2C}$R antagonist, RS 102221 (1 µM), or with application of 5-HT (10 µM). Shaded bars represent time points of values baseline, RS 102221, and after application of 5-HT represented in C.

(B) Averaged firing of baseline (aCSF, gray), RS 102221 (orange), or RS with 5-HT (purple). (ANOVA Treatment: $F_{(1.089, 6.533)} = 2.524$, $p = 0.1593$, n/N = 7/4).

(C) Example electrophysiological traces during application of RS 102221 (left), 5-HT and RS 102221 (middle) and 15 min after 5-HT + RS 102221 washout (right).

(D) Application of WAY 161503 (30 µM) on evoked firing rate of VTA dopamine neurons.

(E) Averaged firing of baseline (aCSF, gray), and WAY 161503 (purple). ($t(5) = 4.828$, $p = 0.0048$, n/N = 6/2).

(F) Example electrophysiological traces during baseline (left), WAY 161503 (middle) and 15 min after WAY 161503 washout (right).

** $p < 0.01$

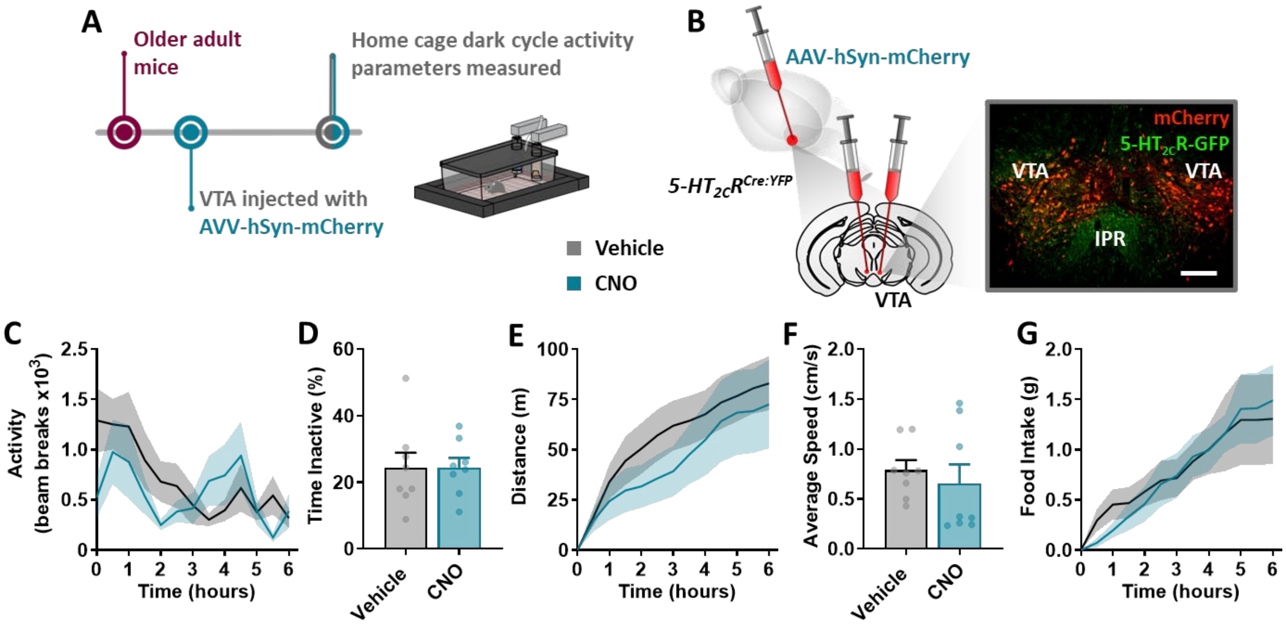


**Figure S4 pertaining to Figure 6. Chemogenetic 5-HT$_{2C}$R$^{VTA}$:mCherry control mice show no changes in physical activity.**

(A) Schematic detailing study design, stereotaxic injection of AAV-hSyn-mCherry control into the VTA (5-HT$_{2C}$R$^{VTA}$:mCherry, in *5-HT$_{2C}$R$^{Cre:YFP}$* mice (12-18 months old, n=8), and assessment with designer drug clozapine-n-oxide (CNO, 1 mg/kg, i.p.) compared with vehicle (saline) treatment.

(B) Schematic detailing AAV-hSyn-mCherry injection site and immunohistochemistry (IF) image showing injection site (red) and co-expression (yellow) with 5-HT$_{2C}$R (green) in the VTA. Scale bar = 200 µm. Ventral tegmental area, VTA; interpeduncular nucleus, IPR. No treatment differences were found including:

(C) total activity in beam breaks (RM ANOVA Treatment: $F_{(1,7)} = 0.9323$, p = 0.3664);

(D) time spent inactive (t(7) = 0.01192, p = 0.9908);

(E) cumulative distance travelled (RM ANOVA Treatment: $F_{(1,7)} = 0.6945$, p = 0.4322);

(F) average speed (t(7) = 0.9554, p = 0.3712);

(G) cumulative food intake (RM ANOVA Treatment: $F_{(1,7)} = 0.01987$, p = 0.8919).

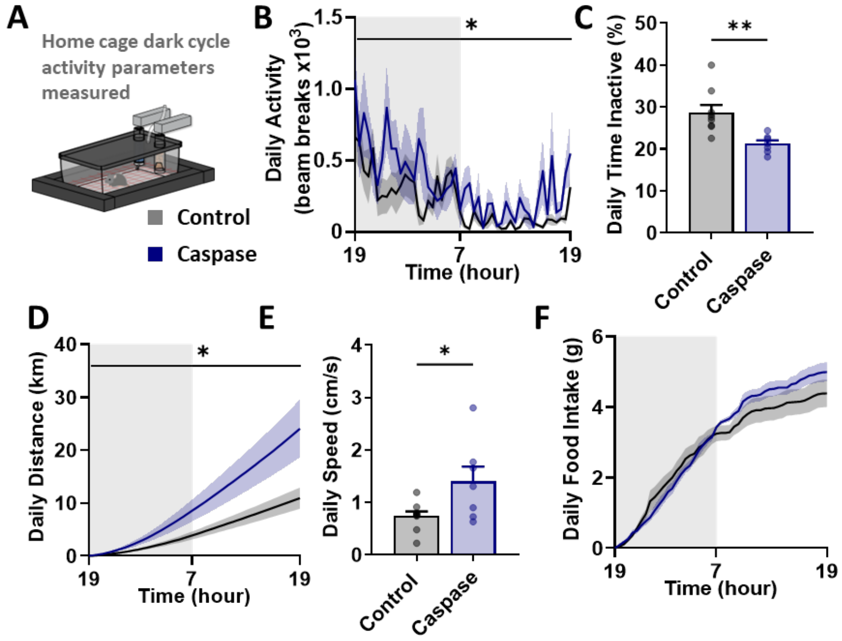


**Figure S5 pertaining to Figure 7. 5-HT$_{2C}$R$^{VTA}$ neuron ablation restores a youthful activity profile in older adult mice over 24 hours.**

(A) Schematic detailing study design and stereotaxic injection of AAV5-Flex-taCasp3-Tevp (n=7) or control (n=9) AAV8-mCherry into the VTA in older adult 5-HT2CRCre:YFP mice (12-18 months old).

(B) Total activity in beam breaks in control and caspase treated mice (RM ANOVA Treatment: $F_{(1,14)}$ = 5.936, p = 0.0288).

(C) Time spent inactive in control and caspase treated mice (t(14) = 3.571, p = 0.0031).

(D) Cumulative distance travelled in control and caspase treated mice (RM ANOVA Treatment: $F_{(1,14)}$ = 5.637, p = 0.0.324).

(E) Average speed in control and caspase treated mice (t(14) = 2.438, p = 0.0287).

(F) Cumulative food intake in control and caspase treated mice (RM ANOVA Treatment: $F_{(1,14)}$ = 0.3448, p = 0.5664).

*p<0.05, **p<0.01.
