## Supplementary material for "Brain serotonin circuit reverses decline in physical activity with age": Table 1

| Variable | Age Group 17-30 (n=45) | 50-70 (n=29) |
|---|---|---|
| **Average Age (yrs)** | **23.29** | **65.69******** |
| StdDev (yrs) | 4.60 | 3.67 |
| **Average Height (m)2** | **1.69** | **1.64*** |
| StdDev (m)2 | 0.10 | 0.08 |
| **Average Body Weight (kg)** | **66.78** | **67.30** |
| StdDev (kg) | 12.98 | 12.48 |
| **Average Waist Circumference (tape, cm)** | **77.41** | **80.23** |
| StdDev (cm) | 11.63 | 13.20 |
| **Average Trunk Fat (%)** | **26.36** | **35.50 **** |
| StdDev (%) | 10.92 | 11.19 |
| **Average Visceral Fat (%)** | **6.82** | **9.75 **** |
| StdDev (%) | 4.21 | 4.58 |
| **Average Percentage of Time at (%)** | | |
| Very Slow Cadance (0-30 steps/min) | **93.84** | **95.24 ***** |
| StdDev (%) | **2.24** | **2.77** |
| Slow Cadance (30-60 steps/min) | **2.18** | **2.27** |
| StdDev (%) | **1.06** | **1.39** |
| Moderate Cadance (60-120 steps/min) | **1.16** | **0.879 *** |
| StdDev (%) | **0.48** | **0.67** |
| High Cadance (90-120 steps/min) | **1.77** | **1.086 **** |
| StdDev (%) | **0.92** | **0.82** |
| Very High Cadance (120-240 steps/min) | **1.08** | **0.518 *** |
| StdDev (%) | **1.32** | **0.63** |
